# *cis*-regulatory architecture of a short-range EGFR organizing center in the *Drosophila melanogaster* leg

**DOI:** 10.1101/366948

**Authors:** Susan Newcomb, Roumen Voutev, Aurelie Jory, Rebecca K. Delker, Matthew Slattery, Richard S. Mann

## Abstract

We characterized the establishment of an Epidermal Growth Factor Receptor (EGFR) organizing center (EOC) during leg development in *Drosophila melanogaster*. Initial EGFR activation occurs in the center of leg discs by expression of the EGFR ligand Vn and the EGFR ligand-processing protease Rho, each through single enhancers, *vnE* and *rhoE*, that integrate inputs from Wg, Dpp, Dll and Sp1. Deletion of *vnE* and *rhoE* eliminates *vn* and *rho* expression in the center of the leg imaginal discs, respectively. Animals with deletions of both *vnE* and *rhoE* (but not individually) show distal but not medial leg truncations, suggesting that the distal source of EGFR ligands acts at short-range to only specify distal-most fates, and that multiple additional ‘ring’ enhancers are responsible for medial fates. Further, based on the *cis*-regulatory logic of *vnE* and *rhoE* we identified many additional leg enhancers, suggesting that this logic is broadly used by many genes during *Drosophila* limb development.

**Author Summary:** The EGFR signaling pathway plays a major role in innumerable developmental processes in all animals and its deregulation leads to different types of cancer, as well as many other developmental diseases in humans. Here we explored the integration of inputs from the Wnt- and TGF-beta signaling pathways and the leg-specifying transcription factors Distal-less and Sp1 at enhancer elements of EGFR ligands. These enhancers trigger a specific EGFR-dependent developmental output in the fly leg that is limited to specifying distal-most fates. Our findings suggest that activation of the EGFR pathway during fly leg development occurs through the activation of multiple EGFR ligand enhancers that are active at different positions along the proximo-distal axis. Similar enhancer elements are likely to control EGFR activation in humans as well. Such DNA elements might be ‘hot spots’ that cause formation of EGFR-dependent tumors if mutations in them occur. Thus, understanding the molecular characteristics of such DNA elements could facilitate the detection and treatment of cancer.

## Introduction

*cis*-regulatory modules (CRMs) are critical for the development and evolution of all organisms. CRMs integrate the information that a single cell or group of cells receives and, in response, trigger changes in cellular and tissue fate specification [reviewed in 1]. The *Drosophila melanogaster* leg imaginal disc, which gives rise to the entire leg and ventral body wall of the adult fly, provides an attractive model system for studying the molecular mechanisms of cellular fate integration at the CRM level and the consequent execution of developmental programs that pattern an entire appendage [reviewed in 2]. Leg imaginal discs are initially specified early during embryonic development through the activation of the transcription factors Distal-less (Dll) and Sp1 in distinct groups of cells in each thoracic segment [3-5]. These groups of cells segregate from the embryonic ectoderm to become the leg imaginal discs, the precursors of the adult legs and ventral thorax. During larval stages, the leg discs proliferate, and defined expression domains of the signaling molecules Wg (ventrally expressed) and Dpp (dorsally expressed) activate *Dll* through the *DllLT* CRM in the center of the leg imaginal disc, where the *wg* and *dpp* expression patterns abut each other (Figure 1 A) [6-9]. Slightly later in development, medial leg fates are established by the feed-forward activation of *dachshund* (*dac*) by Dll through the *dacRE* CRM [2, 10]. During subsequent growth of the leg disc, partially overlapping *Dll* and *dac* expression domains are maintained by autoregulation. These *Dll* and *dac* expression domains, together with the most proximal domain marked by *homothorax* (*hth*) expression, create a rudimentary proximal-distal (PD) axis [2] (Figure 1 A).

**Figure 1.**
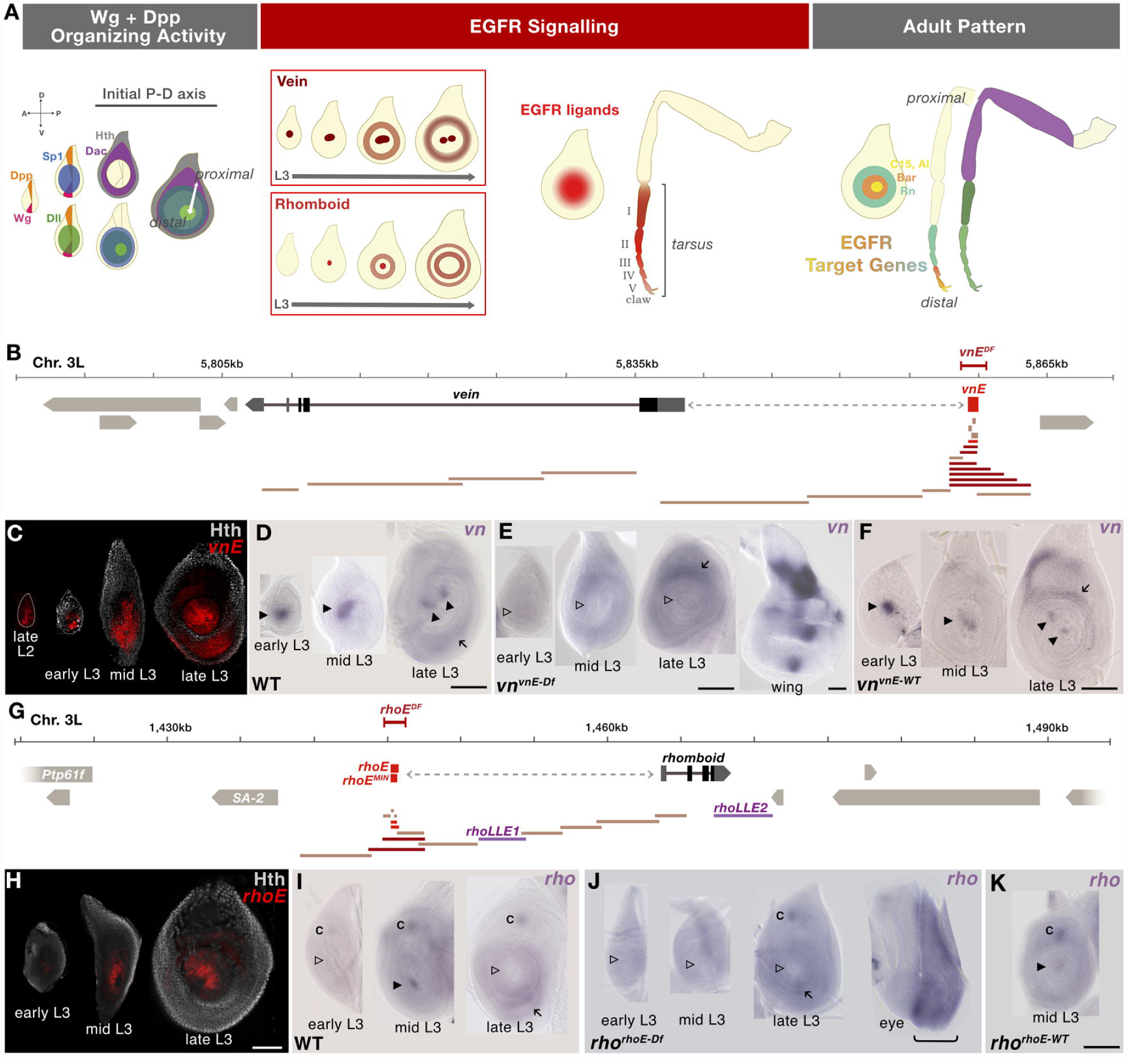
EOC generation in third instar leg discs. (A) Schematic representation of the establishment of an initial PD axis and EGFR signaling events in the center of leg discs (accuracy in the number of depicted rings is not aimed). (B) Schematic representation of the *vn* genomic locus on chromosome 3L; enhancer bashing fragments for identification of *vnE* represented in tan did not drive expression in leg discs, dark red drove expression in leg discs and bright red designates the minimal enhancer used for further analysis; the *vn^vnE-Df^* CRISPR deletion is represented by red bracketed bar. (C) Time-course analysis of the expression pattern of *vnE-lacZ* reporter gene. In young discs, expression is limited to the EOC, while in late L3 additional expression appears in medial rings. (D-F) *In situ* analysis of *vn* expression in 3rd instar leg discs with genotypes: WT (D), *vn^vnE-Df^* (E; a wing disc from the same genotype serves as a positive control). *vn^vnE-WT^*, in which RMCE was used to re-introduce the wild type CRM (F). Arrowheads indicate the presence (filled) or absence (open) of EOC *vn* expression, arrows indicate non-EOC medial expression. (G) Schematic representation of the *rho* genomic locus on chromosome 3L; enhancer bashing fragments for identification of *rhoE* represented in tan did not drive expression in leg discs, dark red drove expression in leg discs and bright red designates the minimal enhancers used for further analysis; *rhoE^MIN^* was only used for enhancer mutagenesis; *rho^rhoE-Df^* CRISPR deletion is represented by the red bracketed bar. (H) Time-course analysis of the expression pattern of *rhoE-lacZ* reporter gene. In young discs, expression is limited to the EOC, while in late L3 additional expression appears in medial rings. (I-K) In situ analysis of the *rho* expression pattern in 3rd instar leg disc with genotype: WT (I), *rho^rhoE-Df^* (J, an eye disc from the same genotype serves as a positive control), *rho^rhoE-WT^*, in which RMCE was used to re-introduce the wild type CRM (K). Arrowheads indicate presence (filled) or absence (open) of EOC *rho* expression, arrows indicate medial expression and “C” indicates chordotonal organ precursor expression. Scale bar = 100μm in all figures.

The initial PD axis defined by Dll, Dac, and Hth is further refined by an additional signaling cascade mediated by the Epidermal Growth Factor Receptor (EGFR) pathway [11, 12]. Like *Dll*, the EGFR pathway is initially triggered by Wg and Dpp, which activate two types of EGFR ligands in the center of the leg disc [11, 12]. One is the neuregulin-related ligand Vein (Vn) and the second is the TGF-!-like ligand Spitz (Spi), which requires metalloproteases of the Rhomboid (Rho) family for processing and secretion [reviewed in 13]. The local activation of *vn* and *rho* family members in the center of the leg disc creates an EGFR organizing center (EOC), a local source of secreted Vn and Spi that activate EGFR signaling in neighboring cells. EGFR signaling in turn results in the activation of a series of downstream target genes that are expressed in nested concentric domains that pattern the future tarsus, the distal-most region of the adult leg [11, 12, 14, 15] (Figure 1 A).

The mechanism by which EGFR signaling patterns the distal leg is not fully understood. One model suggests that EGFR ligands, produced in the EOC, function as morphogens, acting on neighboring cells to generate distinct transcriptional outputs in a concentration-dependent manner. Consistent with this idea is the observation that gradually reducing EGFR activity by raising flies carrying a temperature-sensitive *Egfr* allele (*Egfr^tsla^*) at increasing temperatures results in gradually more severe leg truncations [11]. However, although consistent with a morphogen model, this result is complicated by the fact that in addition to the EOC, there are other sources of EGFR ligands expressed in rings that appear later in leg development [14]. This additional EGFR activation would also be compromised in *Egfr^tsla^* experiments, leaving open the question of the degree to which tarsal PD patterning is due solely to EOC activity. An alternative model posits that the activation of EGFR in the center of the leg disc triggers only local transcriptional outputs, and that alternative sources of EGFR ligands, in combination with indirect transcriptional cascades, are responsible for specifying fates that are further from the EOC.

The EOC/morphogen model predicts that eliminating the production of EGFR ligands from the EOC will have long-range consequences. In contrast, if alternative, non-EOC sources of EGFR ligands play a role in leg patterning, eliminating only the EOC would produce only local defects in distal leg patterning. To distinguish between these models, we searched for CRMs responsible for the expression of EGFR ligands and ligand-processing proteases in the EOC, with the idea that we could specifically eliminate EOC expression by deleting these CRMs. We identified EOC CRMs for *vn* and *rho* (*vnE* and *rhoE*, respectively) and showed that they are necessary for EOC expression of these genes, respectively. However, although EOC expression is eliminated, simultaneous deletion of these CRMs causes only local PD patterning defects and tarsal truncations comparable to mild *Egfr* perturbations in the distal tarsus. These results suggest that the EOC is required for activating local EGFR responses in the center of the leg disc, implying that other sources of EGFR ligands, controlled by non-EOC CRMs, further elaborate the tarsal PD pattern. Finally, we also performed rigorous genetic and biochemical analysis of the *vn* and *rho* EOC CRMs, and used the discovered regulatory logic to predict additional CRMs, many of which are active in the *Drosophila* leg. Together, these data reveal a common regulatory logic for gene activation in the distal leg that is used by many genes, in addition to *vn* and *rho*.

## Results

### Identification and genomic manipulation of the *vn* and *rho* EOC enhancers

To understand the molecular mechanism by which the EGFR signaling pathway is activated in the center of leg imaginal discs during larval stages, we searched for leg disc enhancer elements controlling the expression of EGFR ligands and ligand-processing proteases implicated to function in this process [11, 14]. We scanned the genomic regions of *vn* and *rho* using *in vivo lacZ* reporter assays (Figure 1 B, G and Table S1) and defined minimal enhancers (*vnE* – 654 bp and *rhoE* – 544 bp) that recapitulate the expression pattern of these genes in the center of leg discs during development (Figure 1 C, H), as well as in the serially homologous antennal discs (Figure S1 A, B). The *vnE*- and *rhoE*-*lacZ* transgenes exhibited earlier expression (starting at ~71h PEL for *vnE* and ~82h PEL for *rhoE*; Figure 1 C, H) than detected by *in situ* for *vn* and *rho* (Figure 1 D, I), perhaps because of the greater sensitivity of the anti-ßgal staining, and suggest that the genes might be expressed earlier than previously thought [11, 12, 14, 15]. Our search for leg disc enhancers across the *vn* locus uncovered only *vnE*, while in *rho* we identified two additional *rho* leg disc enhancers (*rhoLLE1* and *rhoLLE2* (Figure 1 G, LLE stands for ‘late leg enhancer’) that drive expression in ring patterns starting in mid-third instar leg discs (90-92h PEL) (Figure S1 C, D). Although these enhancers do not participate in EOC formation, they are active at later developmental stages and drive expression in medial/proximal ring patterns and are thus likely to be additional sources of EGFR activity (Figure S1 C, D).

We also re-examined the expression pattern of additional EGFR ligands and proteases using enhancer-reporter assays (Figure S1 I, L; Table S1), *in situ* hybridization (Figure S1 F, J, M, O; Table S2) and available enhancer trap lines (Figure S1 G) and found that *roughoid* (*ru*) (as previously reported [11]) and *spitz* (*spi*) (Figure S1 G, J), but not *Keren* (*Krn*) or *gurken* (*grk*) (Figure S1 M, O), were expressed in leg discs during third larval instar. Curiously, *ru* expression was only detected by an enhancer trap (*ru^inga^-lacZ*) and by a newly identified enhancer, *ruLLE*, that recapitulates the *ru^inga^* expression pattern (Figure S1 H) but was not detected by *in situ* hybridization (Figure S1 F) (see also Campbell 2002). *spi* was expressed broadly in leg discs (Figure S1 J), and this pattern was recapitulated by a ~10 kb region that includes its promoter and introns (Figure S1 K). Although there are five additional *rho-*family proteases in *Drosophila* [16], previous genetic analysis suggests that *rho* and *ru* are the most relevant [11, 14]. Further, because *ru* did not show expression in early L3 leg discs (and see below for additional genetic tests), and *spi* expression was ubiquitous, we focused on *vnE* and *rhoE* as the primary CRMs active in the leg disc EOC.

To assess the requirement of the *vnE* and *rhoE* CRMs for *vn* and *rho* expression, we deleted them from their native genomic loci using CRISPR/Cas9-mediated genome editing ([17-19]; see Materials and Methods) and assessed the phenotypes of these alleles (*vn^vnE-Df^* and *rho^rhoE-Df^*). We found that these deficiencies abolished the expression of these genes, respectively, only in the EOC of the legs (Figure 1 E, J). The lack of expression in the enhancer deletion alleles was restored when the wild type enhancers were resupplied in their native genomic positions (Figure 1 F, K). Therefore, we conclude that *vnE* and *rhoE* are necessary and sufficient for *vn* and *rho* expression in the EOC, respectively.

### Genetic analysis of *vn^vnE-Df^* and *rho^rhoE-Df^* mutants

Individually, both *vn^vnE-Df^* and *rho^rhoE-Df^* are viable as homozygotes, exhibit normal leg disc patterning (Figure S2 A, C), and form morphologically normal and functional legs (Figure S2 B, D), consistent with previous reports that *vn* and *rho* single mutants do not affect the leg disc or adult leg pattern [11, 12]. However, when we examined the combined effect of these deficiencies in *rho^rhoE-Df^ vn^vnE-Df^* double mutant flies we found that the expression of EGFR downstream genes *C15* and *aristaless* (*al*) was abolished in these animals (Figure 2 A, B and Figure S2 E, F), and the expression of BarH1/H2, a pair of more proximally expressed PD genes [20], collapsed from a ring pattern to a central circular domain in the leg disc (Figure 2 B). In agreement with the leg disc pattern changes, adult *rho^rhoE-Df^ vn^vnE-Df^* double mutants exhibited distal leg truncations that lack a pretarsus and parts of tarsal segment 5 (Figure 2 N). *rho^rhoE-Df^ vn^vnE-Df^* double mutant flies die in late pupal stages most likely because of an inability to exit the pupal case.

**Figure 2.**
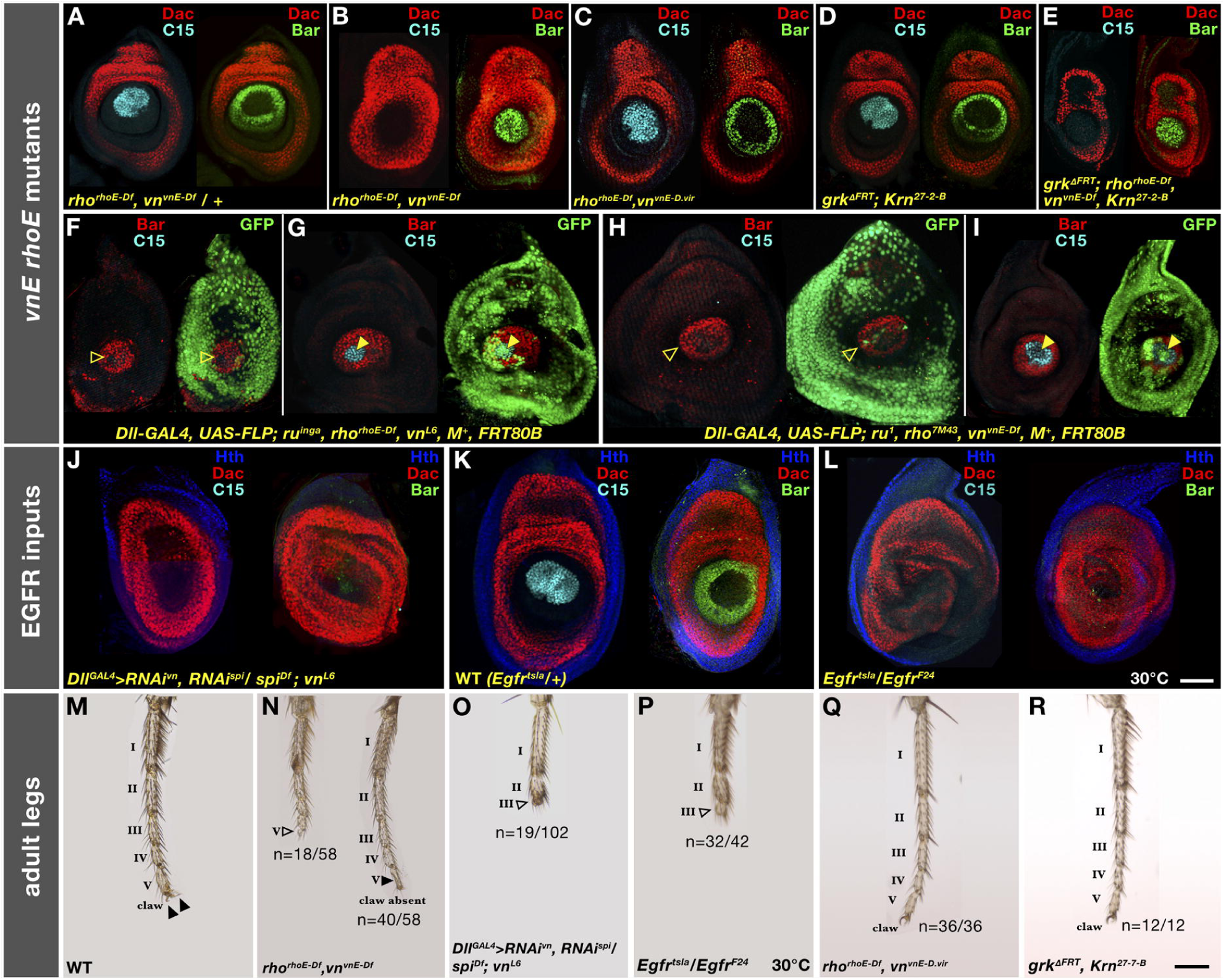
*vnE* and *rhoE* requirement for PD patterning of leg discs and adult legs. (A-L) Effects on distal PD genes (C15 and Bar) in third instar leg discs in: (A) *rho^rhoE-Df^ vn^vnE-Df^ /+*; (B) *rho^rhoE-Df^ vn^vnE-Df^*; (C) *rho^rhoE-Df^ vn^vnE-D.vir^*; (D) *grk*^Δ^*^FRT^; Krn^27-7-B^*; (E) *grk*^Δ^*^FRT^; rho^rhoE-Df^ vn^vnE-Df^ Krn^27-7-B^*; (F,G) *ru^inga^ rho^rhoE-Df^ vn^L6^* mutant clones; (H,I) *ru^1^ rho^7M43^ vn^vnE-Df^* mutant clones; (J) *spi vn* double RNAi; (K) *Egfr^tsla^/+* WT leg disc at restrictive temperature; and (L) *Egfr^tsla^* mutant leg discs at restrictive temperature (M-R) Adult leg morphology of: (M) WT; (N) *rho^rhoE-Df^ vn^vnE-Df^* mutant; (O) *spi vn* double RNAi driven by *Dll-Gal4* (legs with tarsal segments I-II: n=21/102, I-III: n=19/102, I-IV: n=35/102, I-V: n=27/102); (P) *Egfr^tsla^* mutant at restrictive temperature (remaining 10/42 legs less severely truncated); (Q) *rho^rhoE-Df^ vn^vnE-D.vir^*; and (R) *grk*^Δ^*^FRT^; Krn^27-7-B^* mutant. n refers to number of individual legs with a given number of tarsal segments present (even if distal-most segment perturbed). Arrowheads indicate intact (filled) or perturbed (open) tarsal segments.

A sequence comparison between *D. melanogaster* and *D. virilis*, two *Drosophila* species that diverged from each other ~50 million years ago [21], revealed that *vnE* is well conserved (45.8% identity over 0.65 kb) and at a similar location upstream of the *D. virilis vn* transcription start site. In contrast, *rhoE* could not be identified by sequence homology in *D. virilis.* These observations prompted us to ask if the orthologous *D. virilis vnE* (*vnE-D.vir*) could substitute for the function of *D. melanogaster vnE* and rescue the *rho^rhoE-Df^ vn^vnE-Df^* phenotype. We performed the swap of enhancers (see Materials and Methods) and we found that, indeed, the leg imaginal discs of *rho^rhoE-Df^ vn^vnE-D.vir^* flies had normal PD patterning (Figure 2 C) and normal adult legs (Figure 2 Q). This result suggests that the function of *vnE* has been maintained over tens of millions of years and this enhancer element plays a conserved role in limb development.

### Spi and Vn are the relevant EGFR ligands for tarsal leg patterning

Rho is an EGFR ligand-processing metalloprotease that has the potential to cleave the membrane-bound ligands Spi, Krn, and Grk in order to convert them into active secreted forms, while Vn is expressed as a secreted form that does not require Rho function [reviewed in 13]. Although we did not detect any expression of *Krn* and *grk* in leg discs (Figure S1 M, O), this does not exclude the possibility that these genes function in leg disc development at a level of expression below what is detected in our *in situ* hybridization experiments. To address this possibility, we performed genetic experiments and found that the single null mutants *Krn^27-7-B^* [22] and *grk*^Δ^*^FRT^* [23], and the double mutant *grk*^Δ^*^FRT^*; *Krn^27-7-B^*, do not exhibit any leg disc patterning defects (Figure 2 D) or adult leg phenotypes (Figure 2 R). In addition, *rho^rhoE-Df^ vn^vnE-Df^ Krn^27-7-B^* triple mutant (Figure S2 G, H) and *grk*^Δ^*^FRT^*; *rho^rhoE-Df^ vn^vnE-Df^ Krn^27-7-B^* quadruple mutant (Figure 2 E) leg discs had similar defects as *rho^rhoE-Df^ vn^vnE-Df^* double mutants (Figure 2 B), even though the quadruple mutant larvae died at late L3, just before pupation. These results support our conclusion that Krn and Grk are unlikely to be involved in leg development.

The remaining *rho*-dependent EGFR ligand, Spi, is expressed broadly in leg discs (Figure S1 J, K) and is a good candidate for participating in EOC activity under the temporal and spatial control of Rho. To confirm the role of Spi, we used RNAi (see Materials and Methods) to examine the phenotypes of animals depleted for both *spi* and *vn* in leg discs. We found that, indeed, Spi is the EGFR ligand processed by Rho in the center of leg discs, because *spi vn* double RNAi (driven by *Dll-Gal4*) caused loss of expression of the downstream EGFR gene *C15*, and the near elimination of *Bar* expression (Figure 2 J). This phenotype is stronger than any other combination of EGFR pathway components, similar to *Egfr^tsla^* mutants grown at the restrictive temperature of 30°C (Figure 2 L, P). In addition, in animals depleted for *spi* and *vn* using RNAi we observed leg truncations (Figure 2 O) similar to those observed in *Egfr^tsla^* mutants at 30°C (Figure 2 P). Taken together, these results suggest that Vn and Spi are likely the only ligands that activate EGFR signaling during fly leg development.

### Genetic dissection of *rhomboid* and *roughoid* in leg development

The triple *ru^1^ rho^7M43^ vn^L6^* mutant, but not the *rho^7M43^ vn^L6^* double mutant, produces a strong leg truncation phenotype, similar to *Egfr^tsla^* animals grown at 30°C, suggesting that Ru is involved in patterning the adult leg together with Vn and Rho [11]. *vn^L6^* is a nonsense mutation and a null by genetic criteria [24, 25]. *rho^7M43^* is also a null allele [16], although we, as well as previous studies [16], were unable to identify any amino acid changes in the *rho* coding sequence of this allele. *ru^1^* is a nonsense mutation that leads to a premature stop codon after residue 55, prior to the Rhomboid domain, suggesting that it is also a null allele [26]. A potential caveat to this conclusion is that *ru^1^/Df* (including *Dfs ru^PLLb^* and *ru^PLJc^*) results in a stronger ‘rough-eye’ phenotype than the *ru^1^* homozygote, implying that *ru^1^* is a hypomorph [16, 26]. However, *ru,* together with several other genes, is located in the intron of the protein tyrosine phosphatase encoding gene, *Ptp61F*, which plays a role in EGFR/MAPK signaling (Figure S1 E) [27]. Consequently, deficiencies that remove *ru* could also affect M APK/EGFR signaling by reducing *Ptp61F* expression, and could potentially lead to stronger phenotypes compared to the cleaner *ru^1^* allele. Taken together, these observations suggest that *ru^1^* is likely to be a null mutation.

Notably, *rho* and *ru* are physically close to each other on chromosome 3L, with *rhoE* ~55 kb away from the *ru* promoter, raising the possibility that *rhoE* could also regulate *ru* (Figure 1 G and Figure S1 E). To test this possibility, we examined the *lacZ* expression pattern driven by the *ru^inga^* enhancer trap [28] in the background of the homozygous *rho^rhoE-Df^* (see Materials and Methods). We did not detect any effect of *rho^rhoE-Df^* on *ru^inga^-lacZ* expression in leg discs (Figure S2 K, L), suggesting that *ru* is not regulated by *rhoE.*

Because the triple *ru^1^ rho^7M43^ vn^L6^* mutant produces adult leg truncations [11] that are stronger than those observed in our *rho^rhoE-Df^ vn^vnE-Df^* double mutant, we carried out additional experiments to address a potential role for *ru* in leg disc patterning. In the first experiment, instead of examining adult legs we examined *ru^inga^ rho^rhoE-Df^ vn^L6^* triple mutant clones in leg discs (see Materials and Methods). Notably, leg disc patterning in these mutant discs was similar to the pattern observed in the *rho^rhoE-Df^ vn^vnE-Df^* double mutant (Figure 2 F), and even a small patch of WT tissue in the center of the leg disc could restore a normal PD pattern (Figure 2 G). In a second test, we generated *ru^1^ rho^7M43^ vn^vnE-Df^* triple mutant clones and, as in the previous experiment, we observed the loss of C15 and collapse of BarH1 expression (Figure 2 H), similar to the *rho^rhoE-Df^ vn^vnE-Df^* double mutant, and a rescue of C15 expression if some distal cells remain wild type (Figure 2 I).

Together, these results suggest that *ru* does not contribute significantly to EOC activity in the early L3 stage to pattern the L3 imaginal disc. Instead, these results suggest a model in which EOC activity is mediated primarily by *vnE* and *rhoE*, while the later rings of EGFR activation are controlled by a distinct set of enhancers (e.g. *rhoLLE1*, *rhoLLE2*, and *ruLLE*) (Figure S1 C, D, H), and that this second wave of EGFR activity is important for patterning medial regions of the adult leg. In addition, these data suggest that *ru*, and perhaps other *rho*-like family members, plays a role later in leg development through its ring-like expression pattern to ultimately impact adult leg patterning.

### Genetic regulation of *vnE* and *rhoE*

Previous studies have underscored the importance of the Wg and Dpp pathways for EGFR activation in the center of leg discs [11, 12]. Using the *vnE* and *rhoE* enhancer elements, we have been able to address this question in greater detail. We generated mutant clones of *arrow* (*arr*), an obligate co-receptor in Wg signaling, and *Mothers against dpp* (*Mad*), a downstream effector of Dpp signaling, at different time points, and assessed the requirement of these pathways for *vnE* and *rhoE* activation. Both Wg and Dpp pathways are necessary for the initiation of *vnE-lacZ* expression in late L2 larval stage (Figure 3 A, E), while clones made early in L3 stage did not affect *vnE-lacZ* expression (Figure 3 B, F). *rhoE-lacZ* expression was lost when either Wg or Dpp activity was removed during L2 or early L3 (Figure 3 C, G) but became independent of these pathways later in mid-L3 (Figure 3 D, H).

**Figure 3.**
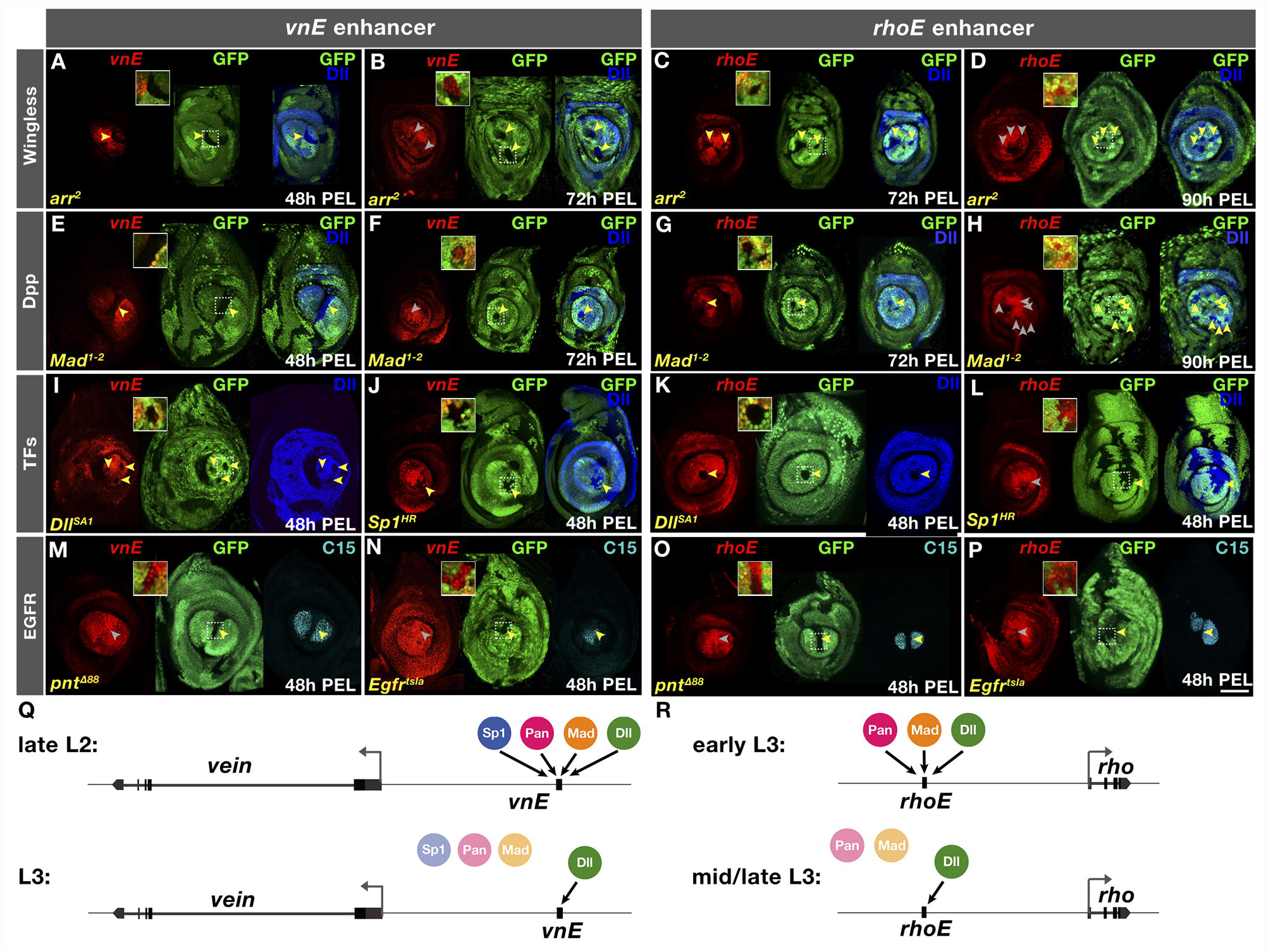
Genetic analysis of inputs into *vnE* and *rhoE*. (A-P) *lacZ* reporter gene expression driven by *vnE* or *rhoE* in mutant clones as indicated. Absence of GFP marks the clone; 2X-magnified insets showcasing specific disc regions (designated with squares) are provided in each case. (A) *vnE* in *arr^2^* clones generated 48h PEL; (B) *vnE* in *arr^2^* clones generated 72h PEL; (C) *rhoE* in *arr^2^* clones generated 72h PEL; (D) *rhoE* in *arr^2^* clones generated 90h PEL; (E) *vnE* in *Mad^1-2^* clones generated 48h PEL; (F) *vnE* in *Mad^1-2^* clones generated 72h PEL; (G) *rhoE* in *Mad^1-2^* clones generated 72h PEL; (H) *rhoE* in *Mad^1-2^* clones generated 90h PEL; (I) *vnE* in *Dll^SA1^* clones generated 48h PEL. (J) *vnE* in *Sp1^HR^* clones generated 48h PEL; (K) *rhoE* in *Dll^SA1^* clones generated 48h PEL; (L) *rhoE* in *Sp1^HR^* clones generated 48h PEL; (M) *vnE* in *pnt*^Δ^*^88^* clones generated 48h PEL; (N) *vnE* in *Egfr^tsla^ Minute+* clones generated 48h PEL; (O) *rhoE* in *pnt*^Δ^*^88^* clones generated 48h PEL; (P) *rhoE* in *Egfr^tsla^ Minute+* clones generated 48h PEL. (Q, R) Schematics summarizing *vnE* and *rhoE* regulation by inputs, respectively.

In addition to Wg and Dpp, at the early larval stages of leg disc development there are two other factors that are crucial for leg specification and growth – the homeodomain transcription factor Distal-less (Dll) [29] and the Zn finger transcription factor Sp1 [4, 5]. *Dll* mutant clones induced at any larval stage abolished *vnE*-lacZ expression (Figure 3I and Figure S3 A). In addition, ectopic expression of *Dll* activated *vnE* not only in leg discs but in other imaginal discs (Figure S3 C, D, E), as long as Wg and Dpp were available in these tissues at the time of clone induction (Figure S3 C, D, E). These results suggest that Dll is required for *vnE* activity. Similarly, *rhoE*-lacZ expression also required Dll at all developmental times (Figure 3K and Figure S3 B).

We also examined the requirement of Sp1 for *vnE* and *rhoE* activation. We found that *vnE* activation requires Sp1, either when the entire animal was mutant or in clones (Figure 3 J and Figure S3 F). This requirement is not mediated by Dll because *Dll* expression remained intact in mutant clones (Figure 3 J) and in leg discs from *Sp1* homozygous animals (Figure S3 F). In contrast, Sp1 was dispensable for *rhoE-lacZ* expression (Figure 3 L and Figure S3 H). In addition, although Sp1 is required for the activation of *vnE* at L2 larval stage (Figure 3 J and Figure S3 F), at the beginning of L3 larval stage Sp1 was no longer required for *vnE* (Figure S3 G). We also assessed if *vnE* and *rhoE* are regulated by *buttonhead* (*btd*), an *Sp1* paralog that is co-expressed with *Sp1* in leg discs [5]. We found that neither EOC enhancer requires *btd* (Figure S3 I, J) and it is unlikely that *rhoE* requires both Sp1 and Btd redundantly since we did not detect Sp1/Btd binding sites or in vivo binding at *rhoE* for Sp1 (see below). Together, these results support a model in which *vnE* activation requires Wg and Dpp together with Dll and Sp1; later, *vnE* activity becomes independent of Wg, Dpp and Sp1, but still requires Dll (Figure 3 Q). Similarly, although the timing differs, *rhoE* requires initial input from Wg, Dpp, and Dll but later only requires Dll for its maintenance (Figure 3 R). The differential onset of expression between the two enhancers might depend on the differential requirement for Sp1.

To investigate if EGFR activity is required for *vnE* and *rhoE*, we examined the expression driven by these CRMs in the background of mutants for EGFR pathway components. *vnE*- and *rhoE*-driven expression was normal in *pnt*^Δ^*^88^* [30] mutant clones or *Egfr^tsla^* [31] mutant clones at the restrictive temperature (Figure 3 M, N, O, P). Capicua (Cic), another downstream component of EGFR [32], is expressed in leg discs (Figure S3 K) but was also not required for *vnE* and *rhoE* activity (Figure S3 L, M).

We next carried out epistasis experiments using the MARCM technique [33] in which we overexpressed one *vnE* or *rhoE* input and removed another. We excluded Sp1 from this analysis because Sp1 sometimes affects Dll expression making results difficult to interpret [5]. For both *vnE* and *rhoE*, we found that while ectopic activation of Dll induced the activity of these enhancers in wildtype tissue (Figure 4 A, E, C, G), in clones compromised for either Wg or Dpp signaling neither *vnE* nor *rhoE* were activated (Figure 4 B, F, D, H). Dll was also unable to induce *vnE*-lacZ expression in ectopic clones in other imaginal discs when Wg and Dpp signaling was compromised (Figure S3 C, D, E). Further, consistent with previous results [6, 7, 34], ectopic Wg and Dpp pathway activity induced *vnE-* and *rhoE-lacZ* expression and created additional EOCs in leg discs when these clones were located close to an endogenous source of Dpp and Wg, respectively (Figure 4 I, M, K, O). However, when these clones were also mutant for *Dll*, these pathways were not able to activate either *vnE* or *rhoE*, and hence EGFR signaling (Figure 4 J, N, L, P).

**Figure 4.**
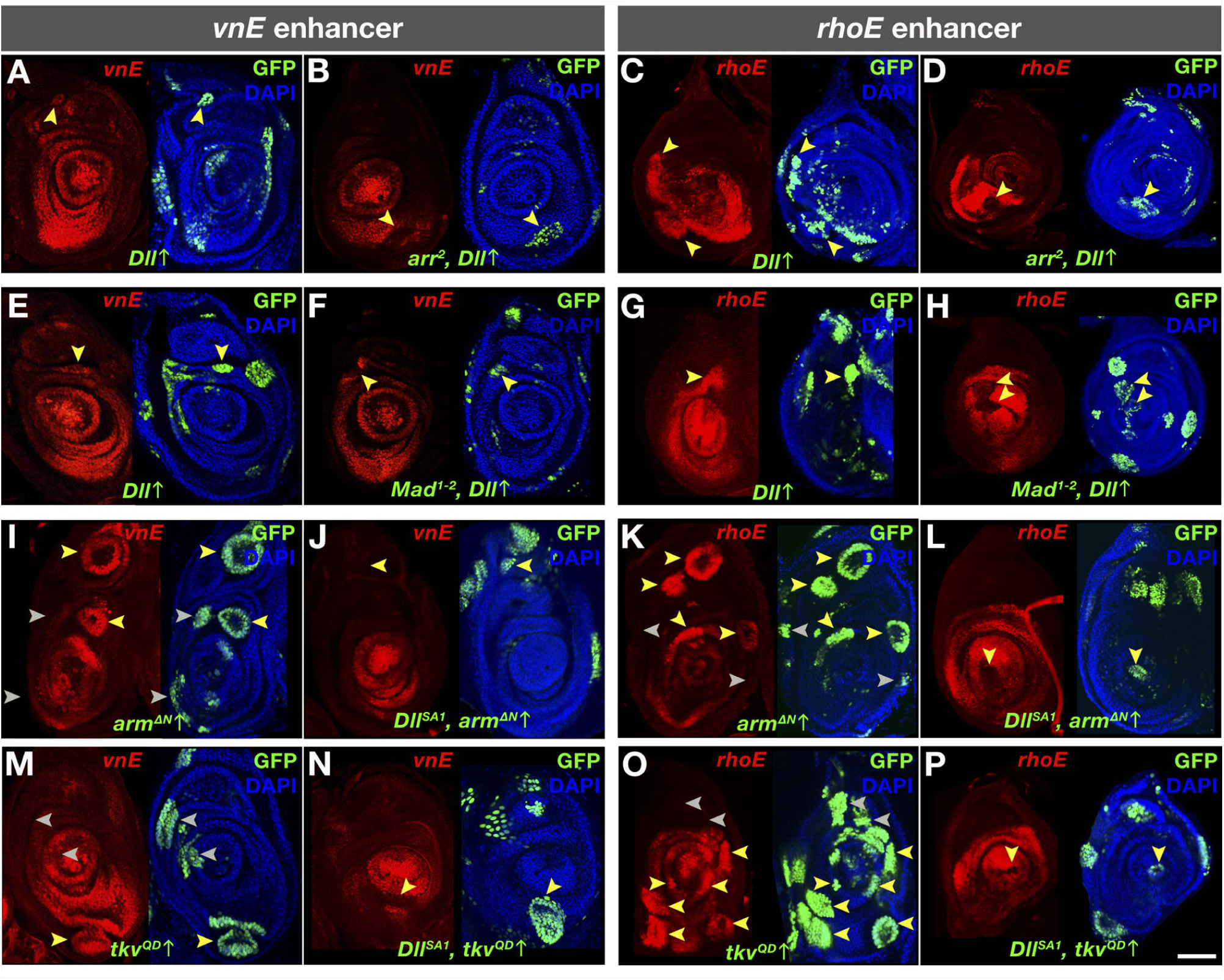
Genetic interactions of inputs into *vnE* and *rhoE*. (A-P) *vnE-* or *rhoE*-directed *lacZ* reporter gene expression in MARCM clones of: (A and C) *Dll* ectopic expression in WT background; (B and D) *Dll* ectopic expression in *arr^2^* mutant background; (E and G) *Dll* ectopic expression in WT background; (F and H) *Dll* ectopic expression in *Mad^1-2^* mutant background; (I and K) *arm*^Δ^*^N^* ectopic expression in WT background; (J and L) *arm*^Δ^*^N^* ectopic expression in *Dll^SA1^* mutant background; (M and O) *tkv^QD^* ectopic expression in WT mutant background; (N and P) *tkv^QD^* ectopic expression in *Dll^SA1^* mutant background

### Dissection of *vnE* and *rhoE* molecular inputs

Our genetic analysis suggests a complex interplay between the signaling pathways Wg and Dpp and the transcription factors Dll and Sp1 on the *vnE* and *rhoE* enhancers. To investigate the configuration of binding sites and the transcription factor grammar of these CRMs, we searched for putative binding motifs using available position weighted matrices (PWMs) [35] and computational methods for identifying consensus Pan (downstream effector of Wg signaling), Mad (downstream effector of Dpp signaling), Dll and Sp1 binding motifs [36]. We performed a comprehensive *in vivo* mutagenesis analysis for both enhancers (Figure 5 A). We mutagenized the enhancer elements by progressively adding (one at a time) mutations (Table S3) in putative binding sites for each transcription factor (Figure 5 A), starting with those that best match consensus binding sites and proceeding to more degenerate binding sites. Because the information from the enhancer bashing experiments (Figure 1 B, G; Table S1) revealed that parts of the enhancers containing multiple sites for each of the TFs can not drive intact expression patterns, we inferred that only having the full set of binding sites gives full expression patterns. Based on the combined analysis between the mutagenesis and the enhancer bashing data we found that there are a large number of binding sites important for *vnE* activation - 14 Pan binding sites, 12 Mad sites, and 11 Dll sites (Figure 5 A, B, D and Figure S4); mutagenesis of subsets of these binding sites leads only to reduction of enhancer-driven expression (Figure S4 A). In contrast, for each TF, there were fewer binding sites important for *rhoE* activation - 4 Pan, 3 Mad and a single Dll binding site (Figure 5 A, C, E). Curiously, in the case of Dll we found 5 additional putative sites in *rhoE* that were not required for enhancer activity in optimal laboratory conditions (Figure S4, Table S3). In general, the identified binding sites for the two enhancers had an additive effect on the expression levels of *vnE* and *rhoE* because partially mutated enhancers drove patchy expression and progressively diminished levels of reporter expression (Figure S4 A, B, C, D). We also confirmed the binding of the TFs involved in *vnE* and *rhoE* regulation by *in vitro* binding assays, suggesting that they act directly to regulate these enhancers (Figure 5 F).

**Figure 5.**
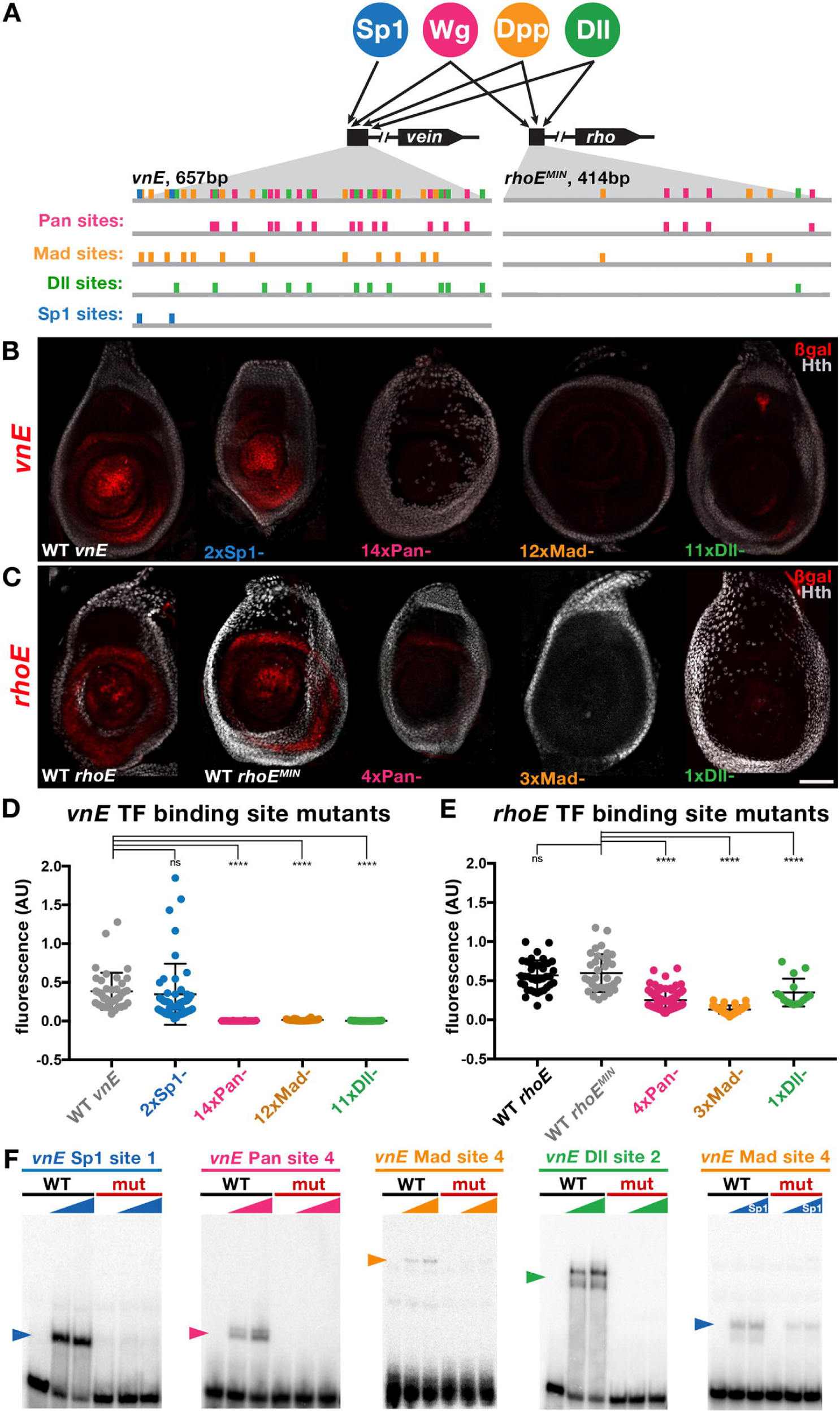
Dissection of Pan, Mad, Dll and Sp1 inputs into *vnE* and *rhoE*. (A) Schematic representation of binding sites in *vnE* and *rhoE*. Putative binding sites for each TF were mutagenized one at a time leading to progressive increase in the number of mutant binding sites until the expression driven by a mutant enhancer was lost. Sites in each TF category that were mutagenized either first or last are not sufficient for full enhancer-driven expression since fragments of *vnE* and *rhoE* (Figure 1 B, G; Table S1) that contain multiple such sites from each TF input cannot drive correct expression pattern. (B) Expression pattern of WT *vnE*-driven expression and mutant *vnE* enhancer-driven expression. (C) Expression pattern of WT *rhoE*-driven expression and mutant *rhoE* enhancer-driven expression. (D) Quantification of WT and mutant *vnE*-driven expression levels in third instar leg discs (WT *vnE* n=41, 2xSp1 n=48, 14xPan n=24, 12xMad n= 32, 11xDll n=49 where n indicates number of leg discs analyzed). For normalization, fluorescence was calculated as a ratio of β-gal:Dll intensity in the center of the leg disc (see Methods for details). (E) Quantification of reduction of WT and mutant *rhoE*-driven expression in third instar leg discs (WT *rhoE* n=39, WT *rhoE^MIN^* n=35, 4xPan n=78, 3xMad n= 41, 1xDll n=15 where n indicates number of leg discs analyzed). For normalization, fluorescence was calculated as a ratio of β-gal:Dll intensity in the center of the leg disc (see Methods for details). (F) EMSA analysis of selected WT vs mutant binding sites.

It is striking that *vnE* contains many more binding sites for each TF compared to *rhoE.* In addition to the differential requirement for *Sp1*, this difference may also contribute to the earlier timing of *vnE* activation compared to *rhoE*, because the larger number of binding sites might render *vnE* more sensitive to lower TF concentrations.

Consistent with the genetic requirement for *Sp1*, we identified two putative Sp1 binding sites in *vnE*. However, when we mutagenized them reporter gene expression was unaltered (Figure 5 B, D). Therefore, we scanned the enhancer by EMSA using overlapping fragments (Table S2) in order to identify additional Sp1 binding sites in an unbiased manner. We found that Sp1 binds with low affinity to some Mad binding sites (Figure 5 F). Because both Sp1 and Mad can bind to some of the same binding sites, loss of *vnE-lacZ* expression when Mad sites are mutated may be a consequence of eliminating all Mad and some of the Sp1 inputs.

Because Sp1 and Dll are co-expressed during leg development, we also scanned all of *vnE* using overlapping oligos (Table S2) to determine if these proteins might bind cooperatively to DNA. For these experiments we used full-length Dll and nearly full-length Sp1 proteins (see Materials and Methods). Although these experiments confirmed Dll binding to its binding sites, we failed to detect any cooperative binding between Dll and Sp1. Taken together, our results suggest that Sp1 regulates *vnE* through two Sp1 binding sites and some shared binding sites with Mad.

### The *vnE* and *rhoE* regulatory logic is widely used among leg CRMs

The *vnE* and *rhoE* regulatory inputs that we discovered here resemble one previously characterized in *DllLT* [9], in that they are all activated by the combinatorial input of Wg, Dpp, Dll and/or Sp1 [5]. These findings prompted us to test if there might be a battery of CRMs that is regulated in the leg disc by these same inputs. To test this idea, we first determined the genome-wide *in vivo* binding profiles of Dll and Sp1 using chromatin immunoprecipitation followed by deep sequencing (ChIP-seq) in third instar leg discs (Figure 6 A). We used either anti-Dll antibody or anti-GFP antibody to ChIP an Sp1-GFP fusion protein expressed from an engineered ~80 kb BAC construct (see Materials and Methods) that drives Sp1-GFP expression identically to *Sp1,* and can rescue an *Sp1* null mutant. Here we focus on genomic loci that show an intersection between 1) Sp1 and Dll binding events, 2) putative Dll, Sp1, Mad and Pan binding sites, and 3) have accessible chromatin as revealed by FAIRE-seq data for leg discs [37]. We found 442 genomic regions that satisfy all six criteria, many of which were close to genes that are expressed in leg discs (Table S4). In addition, two regions correspond to *vnE* and *Dll^M^*, another previously defined CRM of *Dll* (Figure 6). As expected, *rhoE* was not identified because there was no consensus Sp1 binding site in *rhoE*. However, this approach identified a fragment that is within *rhoLLE1* (*rhoLLE1^MIN^*) that, when tested in a reporter gene, drove expression in similar ring pattern as *rhoLLE1* (Figure 6 E).

**Figure 6.**
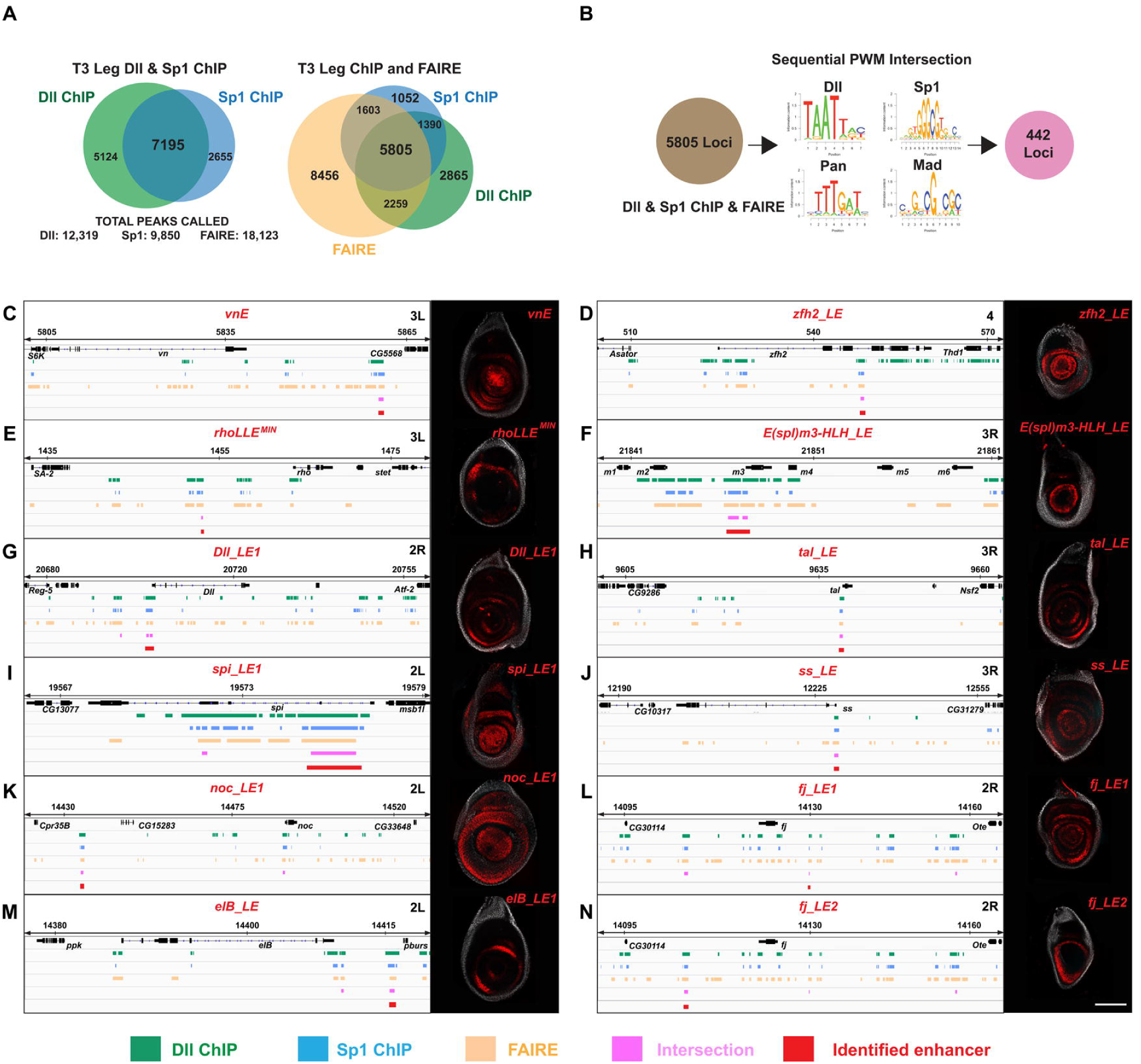
Genome-wide analysis of combinatorial inputs of Dll, Sp1, Wg, and Dpp in leg discs. (A) Venn diagram representing the intersection between Dll ChIP-Seq, Sp1 ChIP-Seq and FAIRE data from third instar leg discs. (B) Schematic representation of bioinformatic intersection between Dll/Sp1 binding events and FAIRE data together with PWMs for Dll, Sp1, Pan and Mad. (C-N) Schematic representation of the binding events at selected genomic loci, the intersections and the expression pattern of tested intersection fragments for: *vnE* (C); *zfh2_LE* (D); *rhoLLE^MIN^* (E); *E(spl)m3-HLH_LE* (F); *Dll_LE1* (G); *tal_LE* (H); *spi_LE1* (I); *ss_LE* (J); *noc_LE1* (K); *fj_LE1* (L); *elB_LE* (M); *fj_LE2* (N).

**Figure 7.**
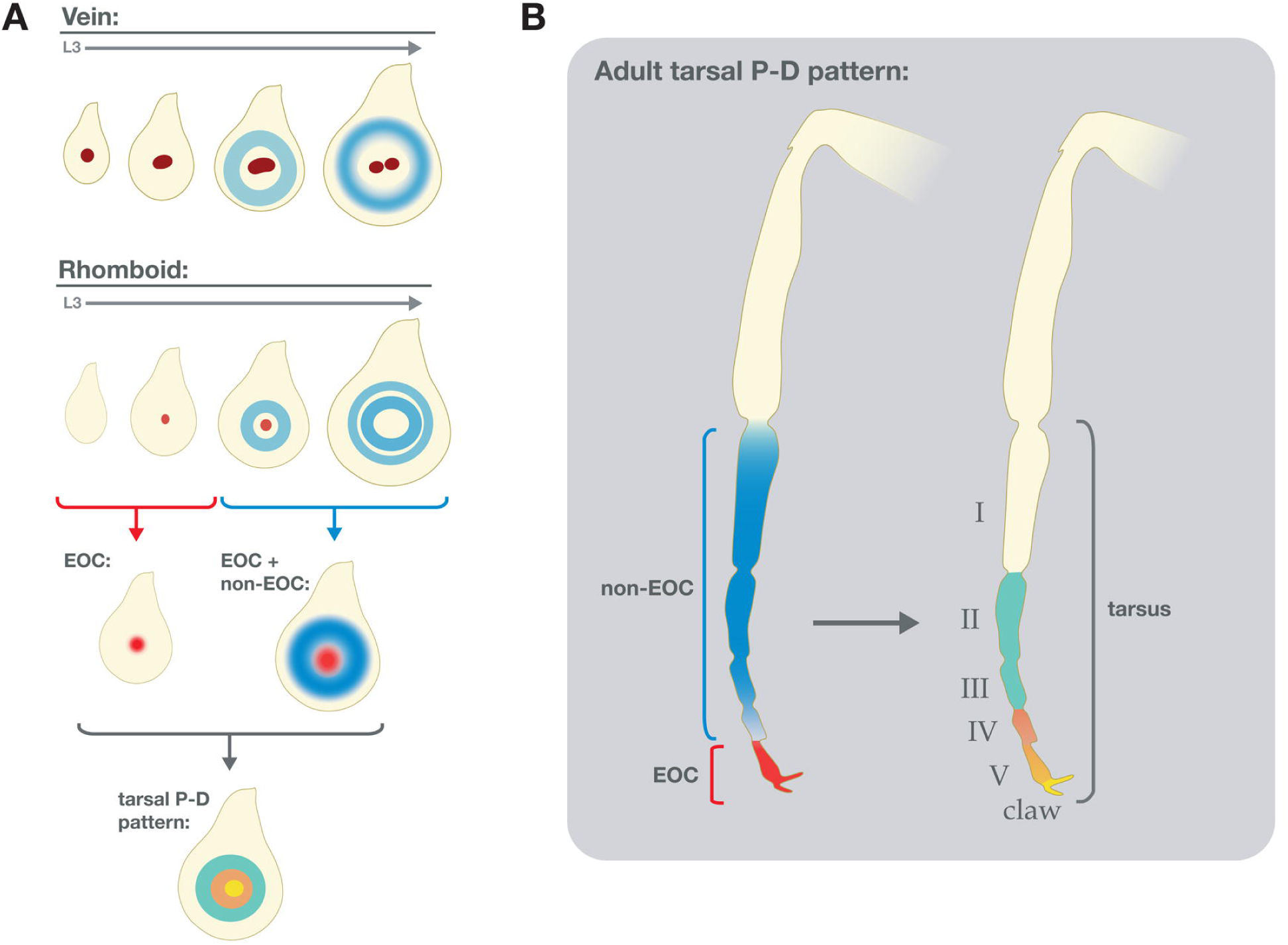
Summary of PD axis patterning by EGFR. Schematic representation of EOC and non EOC sources of EGFR activation along the PD axis in leg discs (A) and adult legs (B).

To validate the larger set of predicted CRMs, we picked 20 additional genomic fragments (23 together with *vnE*, *Dll^M^* and *rhoLL1^MIN^*, ~5% of the total 442 intersections) near 11 genes [*Antennapedia* (*Antp*), *four-jointed* (*fj*), *spitz* (*spi*), *disconnected* (*disco*), *tarsal-less* (*tal*), *spineless* (*ss*), *Zn finger homeodomain 2* (*zfh2*), *elbow B* (*elB*), *no-ocelli* (*noc*), *Enhancer of split m3* (*E(spl)m3-HLH*), and *Distal-less* (*Dll*)]. Using this approach, we discovered at least one leg disc enhancer element with a PD bias for each of the genes we tested (Figure 6 and Figure S5; Table S5), except for *disco*. In some cases (*Antp*, *fj*, *spi, Dll, noc*) multiple fragments generated leg disc expression patterns. Interestingly, we uncovered two leg disc enhancers for the EGFR ligand Spi. Overall, 18 of the 23 tested fragments (78%) are leg enhancers, suggesting that there is a battery of leg disc gene CRMs that drive expression differentially along the leg disc PD axis and are regulated by the direct input of Wg, Dpp, Dll and Sp1.

We also used genome-wide intersection criteria that excluded Sp1 as a factor, thus following the *rhoE* regulatory logic. Not surprisingly, this dataset was much larger (3809 loci), making it difficult to validate experimentally. Nevertheless, it also seems to predict enhancer loci because, in addition to *rhoE*, some of the identified regions corresponded to previously identified leg CRMs such as *Dll DKO* [38], *Dll LL* [39], and enhancer elements identified in genome-wide tiling studies [40].

## Discussion

### Multiple sources of short-range EGFR signaling during fly leg development

The EGFR signaling pathway is widely used in animal development, and is frequently a target in human disease and developmental abnormalities [reviewed in 41]. Yet despite its importance in animal biology, many questions remain about how this pathway functions. Among these questions is whether secreted ligands that activate this pathway can induce distinct cell fates in a concentration-dependent manner. Here, we test this idea by specifically eliminating a single source of EGFR ligands from the center of the *Drosophila* leg imaginal disc, which fate maps to the distal-most region of the adult leg. One plausible scenario is that this single source of secreted EGFR ligands, which we refer to as the EOC, activates distinct gene expression responses at different distances from this source. Alternatively, eliminating ligands secreted from the EOC might only affect gene expression locally, close to or within the EOC. Taken together, our data are most consistent with the second scenario. This conclusion is largely supported by our observations that CRM deletions that eliminate *vn* and *rho* expression from the EOC have mild developmental consequences, both in the L3 leg imaginal discs and adult legs. These phenotypes are significantly weaker than those generated when the entire EGFR pathway is compromised using a temperature sensitive allele of the EGFR receptor. The difference between these two phenotypes is most likely explained by removing only a single source of EGFR ligands in the enhancer deletion experiments versus affecting EGFR signaling throughout the leg disc in the *Egfr^tsla^* experiments. This explanation is further supported by our observation that there are indeed additional CRMs, some of which we define here, that drive EGFR ligand production in more medial ring-like patterns during the L3 stage.

One possible caveat to these conclusions is that there are a total of seven *rho*-like protease genes in the *Drosophila* genome that could, in principle, play a role in distal leg development. We focused on *rho* and *ru*, based on previous results [11, 14] showing that triple *rho ru vn* clones generate severe leg truncations that phenocopy strong *Egfr^tsla^* truncations. In addition, we note that if other *rho* family proteases were active in the EOC, we would not expect to see leg truncations and patterning defects in the leg discs of the *rho^rhoE-Df^ vn^vnE-Df^* double mutant, because those proteases should be able to produce active Spi. These observations suggest that the remaining five *rho*-like protease genes play a minor (or no) role in leg development. However, this conclusion will ultimately benefit from further genetic and expression analysis of these additional *rho*-like genes.

An additional previous observation that contrasts with the suggestion that EOC activity has only a limited role in specifying distal leg fates is the partial rescue of the PD axis when only a small number of distal leg cells were wild type in legs containing large *rho ru vn* clones [11]. However, we note that even in these ‘rescued’ legs, medial defects in PD patterning were apparent. It is also noteworthy that in these earlier experiments, only adult legs were examined. When we repeated the same experiment, but analyzed L3 discs, we found that *rho ru vn* clones generated phenotypes that were very similar to those produced by our double *vnE rhoE* enhancer deletions. Taken together, these observations suggest that timing must be considered in the interpretation of these experiments. When assayed at the late L3 stage, both our enhancer deletion and *rho ru vn* clone experiments argue that EOC activity is limited to specifying only the most distal fates, marked by the expression of *al* and *C15*. Starting in mid L3, and perhaps continuing into pupal development, there are additional sources of EGFR ligands [14] that, when compromised, can affect adult leg morphology. Nevertheless, at least at the L3 stage, these data suggest that EGFR ligands produced from the EOC have a limited and local role in specifying distal leg fates.

### *cis*-regulatory networks during leg development

Integration of inputs from signaling pathways and organ selector genes at CRMs in order to execute distinct developmental programs is a recurrent theme during animal development (reviewed in [42]). Here, we identified two leg EGFR ligand CRMs that integrate the inputs from the Wg and Dpp signaling pathways and the leg selector genes Dll and/or Sp1 in a manner that is very similar to a previously characterized leg enhancer *DllLT* [9]. In addition, when we applied the same regulatory logic to the whole genome, we identified a battery of leg enhancer elements (Figure 6). Interestingly, each of these enhancers drives expression in a specific manner with slightly different timing despite the fact that many of the inputs are shared. It is conceivable that the different expression patterns directed by these enhancers are in part a consequence of additional inputs and/or the difference in the TF binding site grammar. In support of this idea, *vnE* and *rhoE* differ in the number of binding sites for many inputs and *vnE* requires Sp1 while *rhoE* does not. Both of these differences may contribute to the earlier onset of *vnE* expression compared to *rhoE.* The remaining enhancer elements identified in this study direct a plethora of PD-biased leg expression patterns – ranging from ubiquitous, to central and ‘ring’ patterns (Figure 6), which likely integrate inputs in addition to the ones described here. Future studies of these CRMs would help reveal the complex network of regulation that orchestrates leg development in the fruit fly. Such detailed understanding of the *cis*-regulatory architecture of fly leg development would likely give insights into organogenesis and evolution in other animals as well.

### *cis*-regulation of EGFR signaling and cancer

The EGFR signaling pathway has tremendous oncogenic potential and understanding the various mechanisms regulating its activation is not only interesting from the point of view of animal development but also has important practical implications. While the core components of the EGFR pathway have been thoroughly studied because of their potent tumorigenic capability in humans [reviewed in 43], little is known about the transcriptional regulation of EGFR ligands that bind and activate the pathway. The reiterative use of EGFR signaling in many developmental processes implies that different *cis*-regulatory elements are likely utilized by each EGFR ligand in different organs and tissues in order to correctly read the diverse cues in any specific developmental context. It is conceivable that genomic variation in EGFR pathway CRMs might lead to a predisposition to different types of EGFR-dependent tumors in humans, since such CRMs may respond to potent growth-promoting signaling pathways, such as Wnt and BMP.

In this study, we characterized in detail two *Drosophila* EGFR CRMs, *vnE* and *rhoE*, and showed how they integrate the cues from two transcription factors, Dll and Sp1, and two signaling pathways, Wg and Dpp, in order to execute a leg patterning developmental program. Analogous EGFR CRMs are likely to exist in mammals, especially because complex interactions between BMP, Wnt, Shh, multiple Dlx paralogs and other factors, are implicated in the induction of FGF signaling in mammalian limb development. Consistent with this idea, specific single nucleotide polymorphisms (SNPs) in humans in non-coding loci of genes encoding EGFR ligands have been shown to be associated with different types of cancer [44-46]. Such loci may be enhancer elements analogous to *vnE* and *rhoE*. We also note that the regulatory logic uncovered here is likely to be relevant to many CRMs and genes that share spatial and temporal expression programs. Exploiting this regulatory logic in other systems might streamline the identification of enhancer elements that will aid in the discovery of mechanisms that are relevant to EGFR-related human disease and developmental birth defects.

## Materials and Methods

### Drosophila Genetics

The following mutant alleles and enhancer trap alleles were used in this study: *arr^2^, btd^XA^, cic^Q474X^, cic^P[PZ]08482^, dac^p7d23^* (*dac-Gal4*), *Dll^SA1^*, *Dll^em212^* (*Dll-Gal4*), *Egfr^tsla^, Egfr^f24^, grk*^Δ^*^FRT^, Krn^27-7-B^, Mad^1-2^*, *pnt*^Δ^*^88^, rho^7M43^, ru^1^, ru^inga^*, *Sp1^HR^* (shared ahead of publication, [47]), *spi^SC1^, spi^Df(2L)Exel8041^*, *vn^L6^*, *vn^GAL4^*. Transgenic alleles used for *in vivo* clonal ectopic expression of genes were: *UAS-arm*^Δ^*^N^, UAS-tkv^QD^, UAS-Dll.*

To perform RNAi knockdown of *vein* and *spitz* the following strains were used: *UAS*-*vn^RNAi^ (TRiP.HMC04390)/CyO, Dfd:EYFP; UAS-spi^RNAi^ (TRiP.HMS01120)* crossed to either *Dll-GAL4 (Dll^em212^), spi^Df(2L)Exel8041^/ CyO, Dfd-EYFP; vn^L6^/TM6B,* or *spi^Df(2L)Exel8041^/CyO, Dfd-EYFP; vn^GAL4^/TM6B* (*vn^GAL4^* is a null allele [48]). Crosses were raised at 18°C, then shifted to >25°C at the start of L3. For assessment of larval phenotypes, crosses remained at 25°C until fixation and dissection as wandering larvae. For assessment of adult leg phenotypes, crosses were returned to 18°C after 24h until eclosion.

For generation of mutant clones that encompass the entire *Dll*-expressing leg disc region a *yw; Dll-Gal4 (Dll^em212^), UAS-Flp; Ubi-GFP M-y+ FRT80B/C(2L;3R)Tb* strain was crossed to a corresponding FRT80B-containg mutant strain (*ru^inga^ rho^rhoE-Df^ vn^L6^* or *ru^1^ rho^7M43^ vn^vnE-Df^*). For Flp-FRT inducible mitotic recombination and subsequent mosaic clonal analysis fly larvae were heat-shocked at 48h post egg laying (PEL), 72h PEL or 90h PEL and dissected for staining as crawling stage larvae at around 120h PEL. For generation of Flp-FRT mitotic recombination clones, larvae were heat-shocked for 40 minutes at 37°C. Mitotic recombination clones were generated using the following strains: *w hs-Flp^1.22^* Ubi-RFP FRT19A*, yw hs-Flp^1.22^; Ubi-GFP FRT40A /CyO; E/TM6B*, *yw hs-Flp^1.22^; FRT42D Ubi-GFP/CyO; E/TM6B*, *yw hs-Flp^1.22^; E/CyO; Ubi-GFP FRT80B /TM6B*, *yw hs-Flp^1.22^; E/CyO; FRT82B Ubi-GFP/TM6B*, *yw hs-Flp^1.22^; FRT42D M-hs-GFP/CyO; E/TM6B*, *yw hs-Flp^1.22^; E/CyO; Ubi-GFP M-FRT80B/TM6B*. The corresponding strains carrying mutant alleles were used in crosses for generation of mutant clones in the resulting progeny. *E* in these genotypes designates either *vnE-lacZ* or *rhoE-lacZ* inserted in landing sites 51D or 86Fa on chromosome II and III, respectively. To induce GFP expression in larvae marked with *hs-GFP*, an additional heat-shock was given 1 h before dissection for 20 min to 1 hour at 37°C.

The following strains were used for MARCM experiments where *E* designates either *vnE-lacZ* or *rhoE-lacZ* inserted in site 86Fa: *yw hs-Flp^1.22^ tub-Gal4 UAS-GFP; tub-Gal80 FRT40A/CyO; E/TM2*, *yw hs-Flp^1.22^ tub-Gal4 UAS-GFP; FRT42D tub-Gal80/CyO; E/TM2*, *yw; Mad^1-2^ FRT40A; UAS-Dll/C(2L;3R)Tb*, *yw; FRT42D arr^2^; UAS-Dll/C(2L;3R)Tb*, *yw; FRT42D Dll^SA1^; UAS-arm*^Δ^*^N^/C(2L;3R)Tb*, *yw; FRT42D Dll^SA1^; UAS-tkv^QD^/ C(2L;3R)Tb*, *yw; y+ FRT40A; UAS-Dll/C(2L;3R)Tb*, *yw; FRT42D y+; UAS-Dll/C(2L;3R)Tb*, *yw; FRT42D y+; UAS*-*arm*^Δ^*^N^/C(2L;3R)Tb*, *yw; FRT42D y+; UAS-tkv^QD^/ C(2L;3R)Tb*.

For all *in vivo* clonal experiments, at least 20 examples of discs with clones of the correct genotype were examined, which is typical for experiments of this type, and more than one independent experiment was carried out for each tested genotype.

### Plasmids and transgenes

All wildtype and mutagenized *enhancer-reporter* transgenic constructs were made using the *lacZ* reporter vector pRVV54 as an acceptor vector [49]. Coordinates of the genomic fragments PCR-amplified in the enhancer bashing experiments are listed in Table S1 and Table S5. The ΦC31 system was used for transgenesis and plasmids were introduced in landing sites 51D or 86Fa [50].

Site-directed mutagenesis of the *vnE* and *rhoE* enhancers was performed according to the QuikChange II protocol (Agilent Technologies). *vnE* and *rhoE* enhancers were first introduced in pBluescript SK+ vector for site-directed mutagenesis and the resulting mutated enhancers were consequently transferred to pRVV54 for *in vivo* analysis in the fruit fly. Primers used for mutagenizing of putative binding site are listed in Table S3.

Plasmids for recombinant protein production were made by introducing cDNA sequences into pET21 series vectors (Novagen-EMD Millipore) and their derivatives, resulting in C-terminally tagged His proteins. Primers used to generate Dll-His (full-length Dll), Sp1^Zn-finger^-His (only the Zn-finger domains; used for confirming *in vitro* binding to Sp1 sites), Sp1^424AA^-His (used to examine cooperativity with Dll), Mad^MH1^-His (only the MH1 domain) and Pan^HMG^-His (only the HMG domain) vectors are listed in Table S2.

### CRISPR/Cas9 alleles

The *vnE* and *rhoE* CRISPR/Cas9 alleles were generated by using pCFD4 vector for driving gRNA expression [18] and a germline-expressing Cas9 donor strain for plasmid mix injection [19]. The following sequences were used as gRNAs for generation of the *vnE^Df^* allele: CGATTTTAATGCGAAAGCTA and TTTGGCTTTCAACGCTTAAT. The following sequences were used as gRNAs for generation of the *rhoE^Df^* allele: GAGCCGAGGGCACAAATTGA and ATGATGATGATGTATTGCCC. We created a vector containing a cassette with *P3-RFP* [50] and *FRT(F5)-hs-neo-FRT(F5)* selectable markers flanked by minimal inverted ΦC31 [51] attP sites (pRVV613) [52]. This vector was used for insertion of upper and lower homologous arms for generation of donor vectors for creation of platforms for cassette-exchange. Primers used for PCR-amplification of the homologous arms are listed in Table S2. *vnE* and *rhoE* pCFD4-based gRNA vectors (250ng/μl) were co-injected with the corresponding *vnE* and *rhoE* homologous arm donor cassette vectors (500ng/μl) and resulting flies were screened for P3-RFP expression. To generate *rhoE* deletion allele in the background of *ru^inga^*, injections to generate the *rhoE^Df^* were repeated in a *nos-Cas9/CyO; ru^inga^/TM3* strain. Positive fly lines were verified by PCR for correct insertion of the donor cassettes. Deletion alleles without *P3-RFP* were generated through RMCE by injection with an empty multiple cloning site vector containing inverted ΦC31 attB sites (pRVV578) [52]. The *P3-RFP*-containing and -non-containing enhancer deletion alleles exhibited identical expression patterns and phenotypes. The WT *vnE*, *rhoE* and the *D. virilis vnE* enhancers were cloned into pRVV578 and resupplied by RMCE in a similar manner (primers are listed in Table S2).

### Protein assays

Recombinant proteins were expressed in BL21 (DE3) cells (Agilent Technologies) through IPTG induction for 4h. Proteins were subsequently purified through Cobalt chromatography with TALON Metal Affinity Resin (Clontech, #635501). EMSA gels were performed as previously described [53].

### Immunohistochemistry and adult leg analysis

Immunostainings of fly imaginal discs was performed by standard protocol. The following antibodies were used in this study: rabbit anti-β-galactosidase (Cappel), mouse anti-β-galactosidase (Sigma-Aldrich, #G4644), guinea pig anti-Dll [9], rat anti-Sp1, guinea pig anti-Hth [54], mouse-anti-GFP (ThermoFisher Scientific, #A11121), rat anti-C15 [15], rat anti-Al [6], rat anti-BarH1 [55], rabbit anti-BarH1 [56], mouse anti-Dac [57]. AlexaFluor488-, AlexaFluor555-, and AlexaFluor647-conjugated secondary antibodies from ThermoFisher Scientific or Jackson ImmunoResearch Laboratories were used at 1⋮500 dilution.

Adult legs were dissected, mounted, and analyzed by light microscopy. All adults of the relevant genotype that eclosed within an 8-hour period were scored. Roman numerals in the figure legends indicate the tarsal segments present in each phenotypic class (with the distal most segment perturbed). For example, a truncation designated as I-III means that tarsal segments I, II and III were present, with segment III partially defective (e.g. Figure 2 P). n refers to the number of individual legs scored. The number of legs examined for each genotype is reported in the figures and figure legends.

### In situ Hybridization

To generate vectors for *in situ* probes *vn*, *ru*, *spi*, *Krn*, and *grk* DNA sequences were amplified from genomic DNA and *rho* DNA sequence was amplified from cDNA clone (LD06131; DGRC clone #3528) using primers listed in Table S2. DNA fragments were cloned into pBluescript SK+ (Agilent Technologies).

RNA antisense probes were transcribed with either T3 or T7 RNA polymerase (depending on the cDNA sequence orientation in the vectors listed in Table S2) and labeled using DIG UTP mix (Sigma, #11175025910). Sense RNA probes were used as negative controls. *rho* probes were then hydrolized for 30 minutes at 60°C as previously described [58]. Third instar larvae were dissected in cold 1xPBS and fixed for 16h at 4ºC in 4% PFA + 2mM EGTA. *In situ* hybridization was then performed as previously described [58] and signal was developed in BM-Purple AP substrate (Sigma #11442074001) after staining with anti-DIG - AP antibody at a concentration of 1:2000 (Roche #1093274). Multiple (#10) discs were examined for each time point, probe, and genotype.

### Fluorescence Quantification

Mid-third instar larvae carrying wild-type or mutant *vnE-* or *rhoE-lacZ* reporter constructs were raised, fixed, stained and imaged in parallel according to standard immunohistochemical protocols. Average fluorescence was measured for the area within the central/tarsal domain of all unobstructed leg imaginal discs using ImageJ software (http://rsb.info.nih.gov/ij) and reported as the ratio of β-gal:Dll (staining control) in arbitrary units (AU). Ordinary one-way ANOVA adjusted for multiple comparisons (Dunnett’s test) were performed and graphed in Prism software (graphpad.com) to compare wild-type fluorescence to mutant enhancer genotypes where ns = not significant, * = p ≤ 0.0332, ** = p ≤ 0.0021, *** = p ≤ 0.0002 and **** = p<0.0001 (adjusted p-values). n refers to the number of individual leg discs scored. The number of leg discs scored for each genotype is reported in the figure legends.

### Chromatin IPs

Triplicate pools of 100 *yw* and 100 *Sp1-GFP^BAC^* L3 wandering larvae were used to perform independent chromatin IPs as previously described [59]. The *Sp1-GFP^BAC^* is a GFP-tagged Sp1 in BAC clone CH321-64M02 inserted in landing site VK00033 (gift from Dr. Rebecca Spokony). All 6 leg discs from each larva were used as material for each IP. Chromatin from the *yw* larvae pools was immuno-precipitated with goat anti-Dll antibody (sc-15858, Santa Cruz Biotechnology, 1.5 μg/ml for IP) while chromatin from the *Sp1-GFP^BAC^* larvae pools was immuno-precipitated with rabbit anti-GFP antibody (ab290, Abcam, 1⋮300 dilution for IP). DNA from non-immunoprecipitated 10% chromatin input was isolated from each pool as reference control. Both control and immunoprecipitated DNA samples were prepared for Illumina sequencing using the Epicentre Nextera DNA Sample Preparation Kit and sequenced on an Illumina HiSeq 2000 according to the manufacturer’s specifications. Experiments were performed in duplicate and peak calling was based on merged reads for duplicate ChIPs. Sequences were aligned to the Drosophila genome using the Burrows-Wheeler Aligner and ChIP-seq peaks were called using MACSv2 [60, 61]. Peak regions were defined using a p-value cutoff of 1.00e-02, but only those peaks passing a more stringent q-value cutoff of 1.00e-04 were used for further analysis. Datasets generated in this study are available at the Gene Expression Omnibus (GEO): accession number GSE113574, https://www.ncbi.nlm.nih.gov/geo/query/acc.cgi?acc=GSE113574.

### Bioinformatic intersection analysis

PWMs for Dll, Sp1, Pan, and Mad were extracted from The Fly Factor Survey Database using the command *grep* within the MotifDb Bioconductor/R package. To generate BED files containing position information for each of the above PWMs, the *matchPWM* command from the Biostrings Bioconductor/R package was used. In-house code was used to run the command iteratively through the chromosomes (using DM3 build). Only hits above a minimum score of 80% were retained. IGVtools within the Integrative Genomics Viewer (IGV) was used to sort and index the BED files prior to intersection. Intersections of all BED files (derived from PWM analysis and ChIP-seq and FAIRE peak calling analysis) were done using Bedtools2 run locally from the command line. ChIP-seq peaks for Dll and Sp1 were first intersected with the FAIRE peaks. The product of this intersection was then sequentially intersected with each of the PWM files, always returning the peak coordinates from the initial file. The command *intersectBed* was used with options: -wa, -F 1.0, -u. To determine the gene nearest to each of the intersected ChIP peaks, packages within R/Bioconductor were used. The annotation package TxDb.Dmelanogaster.UCSC.dm3.ensGene was downloaded and annotated transcripts extracted. The *distanceToNearest* function was used to find the nearest annotated transcript to each of the ChIP Peaks. In-house R script was then used to generate the table containing the coordinates of the ChIP peaks, as well as the nearest annotated gene (Table S4).

## Acknowledgements

We are grateful to Drs. Myriam Zecca, Yannis Mavromatakis, Gerard Campbell, Gary Struhl, Amanda Simcox, Benny Shilo, Andrew Tomlinson, Carlos Estella, Jessica Treisman, Josepha Steinhauer, Gerardo Jimenez, Rebecca Spokony and Trudi Schupbach for sharing numerous reagents that greatly facilitated this study. We thank members of the Mann lab, Gary Struhl, and Harmen Bussemaker for comments and suggestions during the course of these studies. R.V. is a Leukemia and Lymphoma Society Fellow. This work was supported by NIH grants R35GM118336 and RO1GM058575 awarded to R.S.M.

## Author contributions

RV and RSM conceived this work and wrote the article. RV, SN, and RSM designed the experiments and analyzed the data. RV and SN performed the experiments and made the figures. AJ performed the ChIPs, MS generated sequencing libraries; AJ and MS processed the ChIP-seq data. RKD performed the bioinformatic analyses.

---

**Figure S1.**
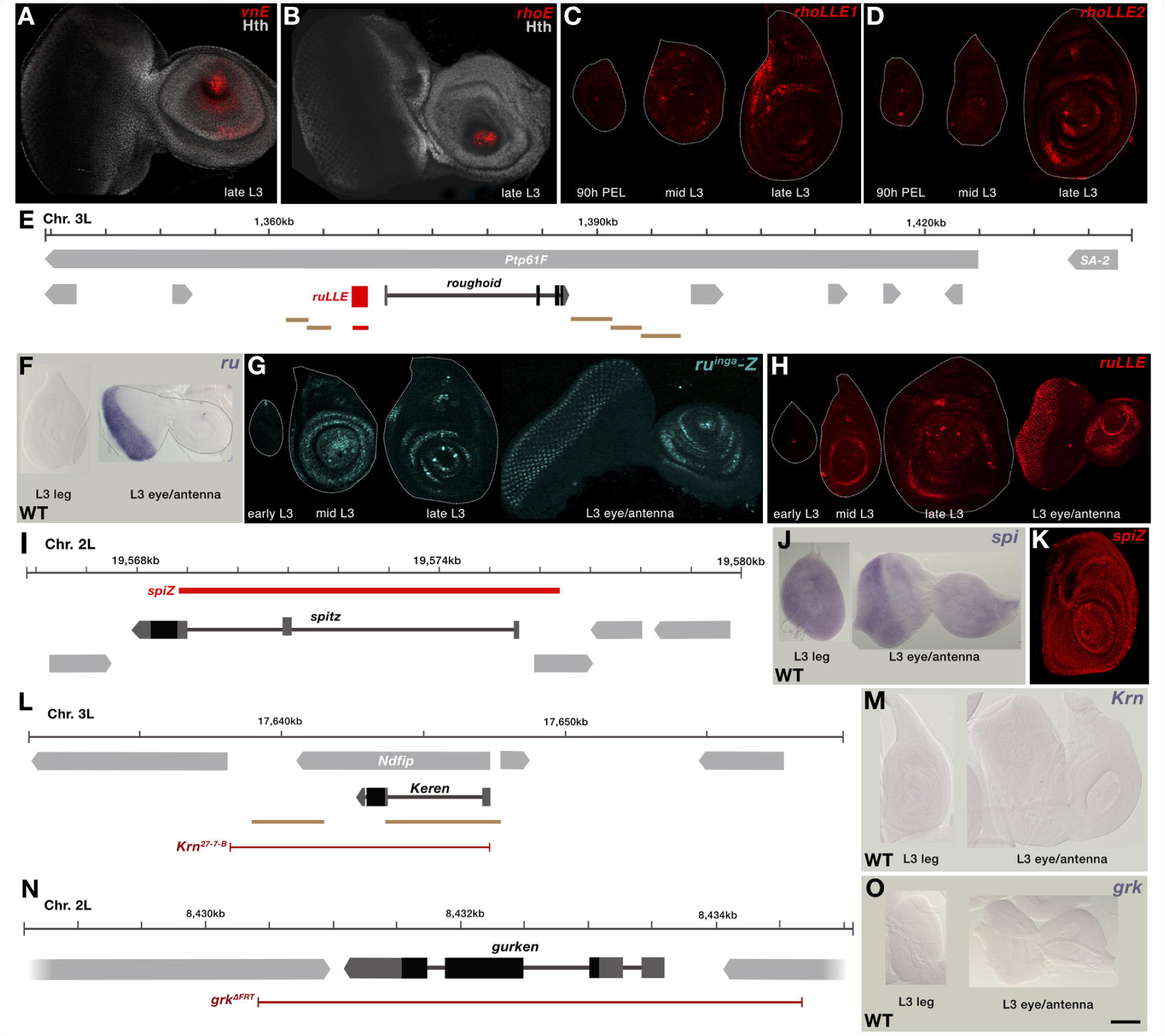
Expression pattern of additional *rho* enhancers, EGFR ligands and ligand-processing proteases. (A-B) Expression pattern of *vnE* (A) and *rhoE* (B) in third instar eye-antennae discs. (C-D) Expression pattern of (C) *rhoLLE1* and (D) *rhoLLE2* throughout leg disc development. (E) Schematic representation of *ru* genomic locus with enhancer bashing results. Fragments represented in tan did not drive expression in leg discs. (F-H) Expression pattern of *ru* from in situ (F), *ru^inga^* (G) and *ruLLE* (H). (I) Schematic representation of *spi* genomic locus with enhancer bashing results. (J-K) Expression pattern of *spi* from in situ (J) and *spi-lacZ* reporter construct (K). (L) Schematic representation of *Krn* genomic locus with enhancer bashing results and *Krn^27-7-B^* mutant. Fragments represented in tan did not drive expression in leg discs. (M) Expression pattern of *Krn* from in situ. (N) Schematic representation of *grk* genomic locus with *grk*^Δ^*^FRT^* mutant. (O) Expression pattern of *grk* from in situ.

**Figure S2.**
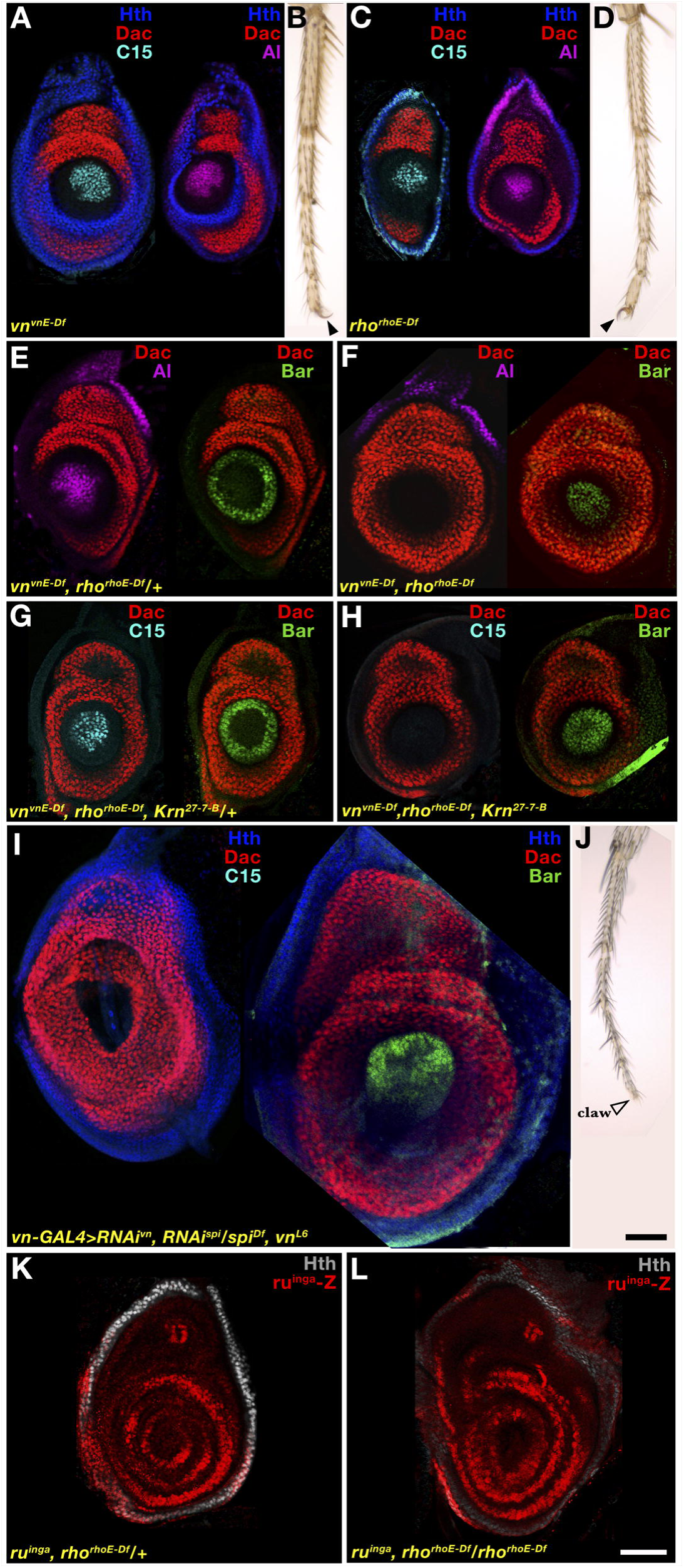
Additional genetic analysis of *vn^vnE-Df^*, *rho^rhoE-Df^* and other EGFR-activating components. (A) Expression pattern of C15 and Al in *vn^vnE-Df^* mutant. (B) Adult leg of *vn^vnE-Df^* mutant. Filled arrowhead indicates intact pretarsal claw. (C) Expression pattern of C15 and Al in *rho^rhoE-Df^* mutant. (D) Adult leg of *rho^rhoE-Df^* mutant. Filled arrowhead indicates intact pretarsal claw. (E) Expression pattern of Al and BarH1 in WT (*rho^rhoE-Df^ vn^vnE-Df^*/+) and (F) *rho^rhoE-Df^ vn^vnE-Df^* double mutant. (G) Expression pattern of C15/Bar/Dac in WT (*rho^rhoE-Df^ vn^vnE-Df^ Krn^27-7-B^*/+) and (H) *rho^rhoE-Df^ vn^vnE-Df^ Krn^27-7-B^* triple mutants. (I-J) *spi vn* double RNAi driven by *vn-GAL4.* Expression pattern of C15 and BarH1 in third instar leg discs (I) and adult leg (J). Open arrowhead indicates absent pretarsal claw). (K-L) Expression pattern of *ru^inga^-lacZ* in *ru^inga^ rho^rhoE-Df^/+* (K) and *ru^inga^ rho^rhoE-Df^*/ *rho^rhoE-Df^* (L) leg imaginal discs.

**Figure S3.**
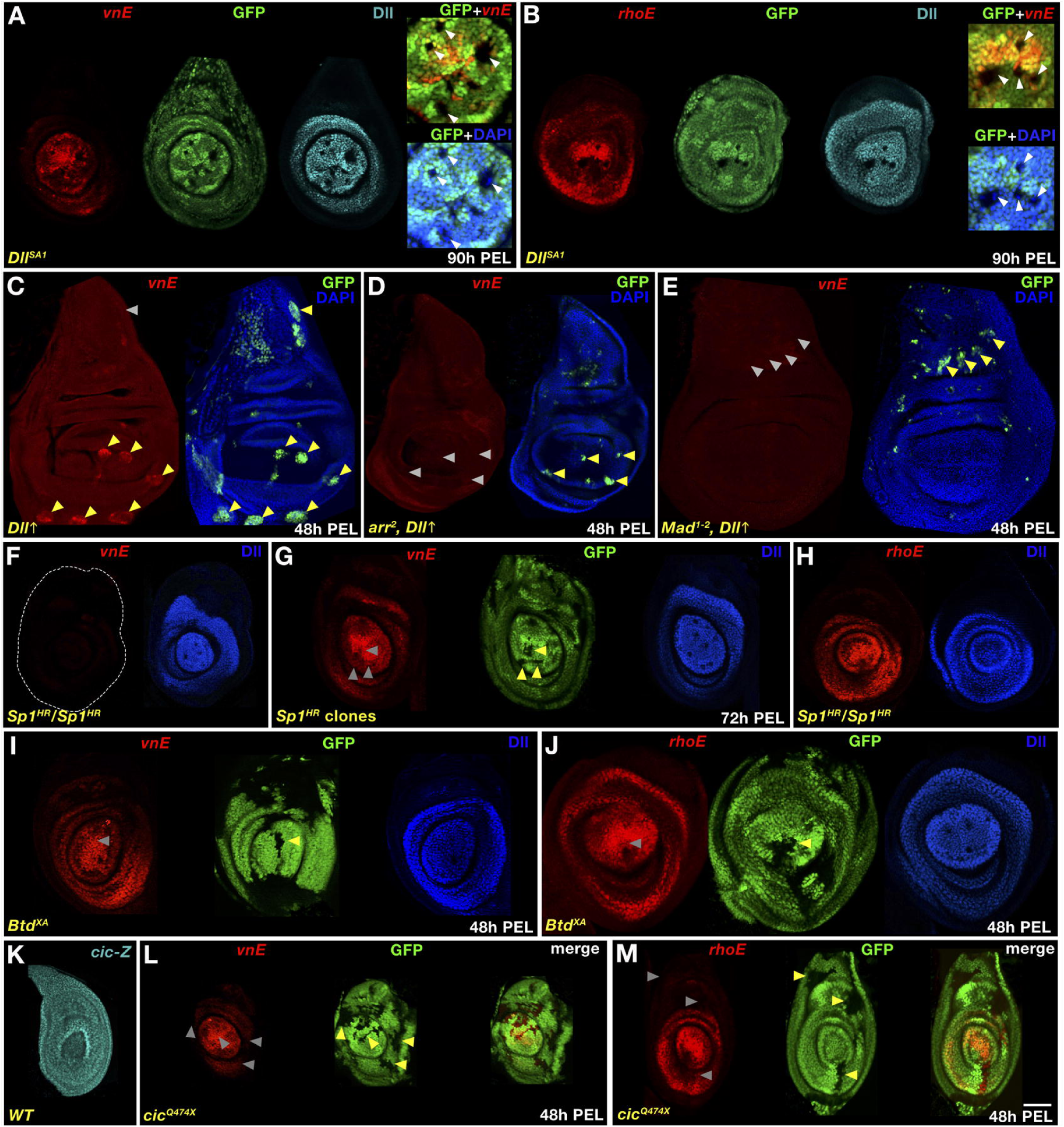
Additional genetic analysis of *vnE* and *rhoE* inputs. (A) *vnE*- or (B) *rhoE*-driven *lacZ* expression in *Dll^SA1^* mutant clones generated at 90h PEL. (C-E) *vnE*-driven *lacZ* expression in wing discs in *Dll* ectopic overexpression clones in WT background (C); *Dll* ectopic overexpression clones in *arr^2^* mutant background (D); *Dll* overexpression clones in *Mad^1-2^* mutant background (E); (C-E) clones were generated at 48h PEL. (F-G) *vnE*-driven *lacZ* expression in leg discs of *Sp1^HR^* mutant animals (F); *Sp1^HR^* mutant clones generated at 72h PEL (G). (H) *rhoE*-driven *lacZ* expression in leg discs of *Sp1^HR^* mutant animals. (I-J) *vnE*- (I) or *rhoE-* (J) driven *lacZ* expression in *btd^XA^* mutant clones generated at 48h PEL. (K) *cic-lacZ* expression in leg discs. (L-M) *vnE*- (L) or *rhoE*- (M) driven *lacZ* expression in leg discs with *cic^Q474X^* mutant clones generated at 48h PEL.

**Figure S4.**
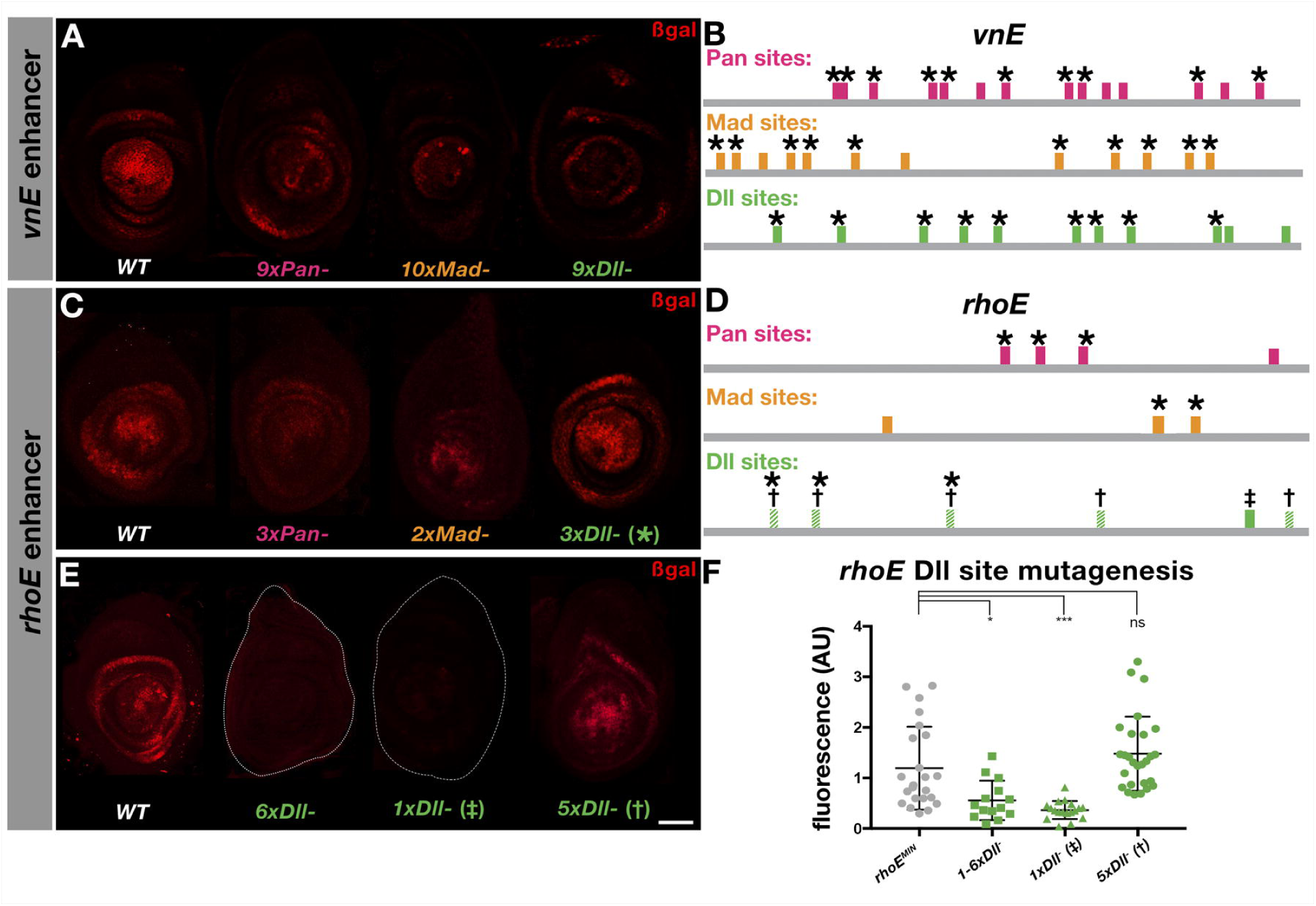
Expression of partially mutant *vnE* and *rhoE* reporter genes. (A, C, E) *vnE*- (A) and *rhoE-* (C, E) driven expression of WT and intermediately mutant CRMs. (B and D) schematic representation of binding sites in *vnE* and *rhoE*, respectively. Mutated sites for the CRM-reporter genes shown in A, C, and E are indicated by the *, † and ‡. (F) Quantification of expression levels; fluorescence was calculated as a ratio of β-gal:Dll intensity in the center of the discs (see Methods for details). WT *rhoE^MIN^* n=23, 6xDll n=14, 1xDll n= 18, 5xDll n=27 where n indicates number of leg discs analyzed.

**Figure S5.**
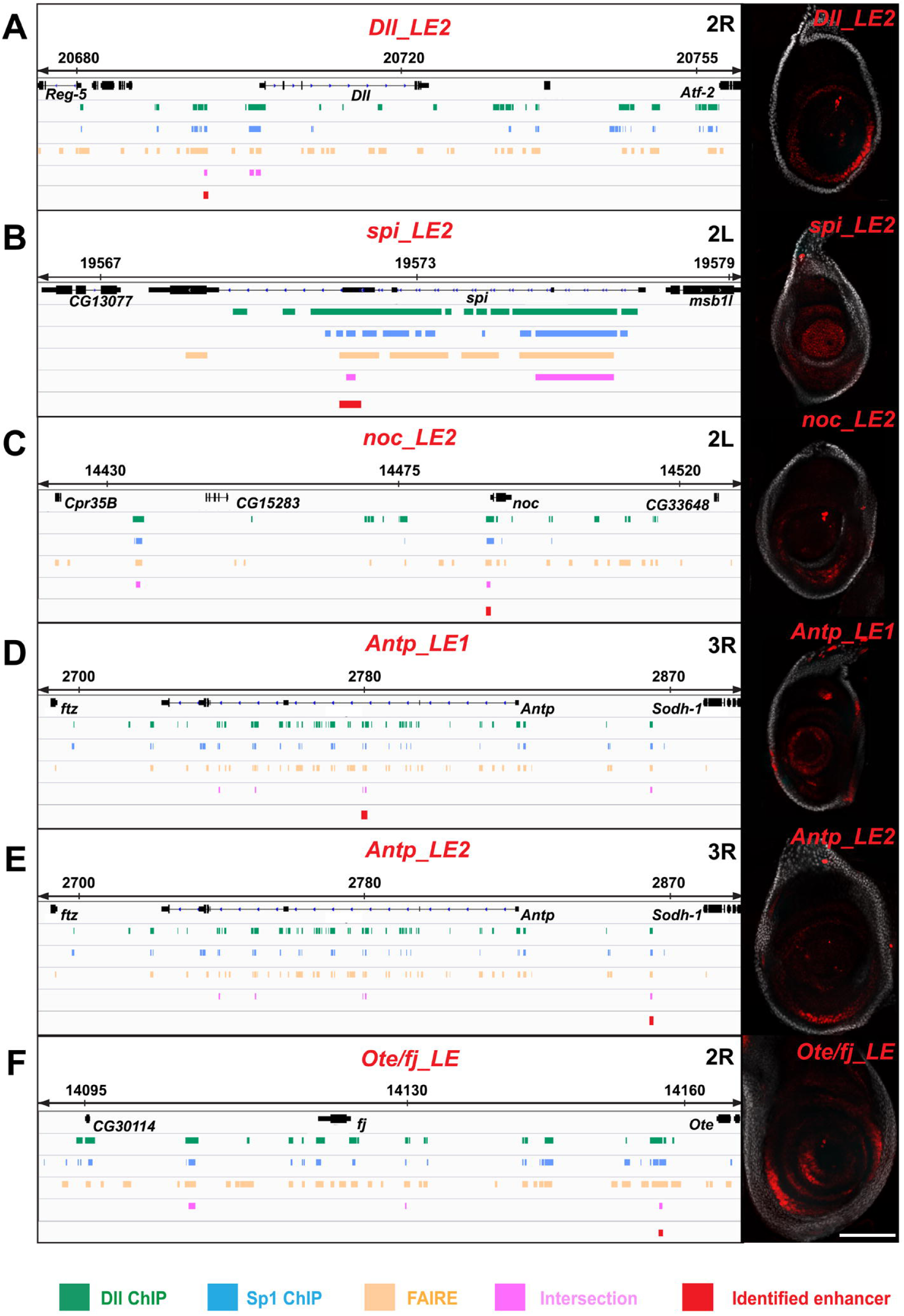
Additional identified enhancers. Schematic representation of identified genomic loci and the expression patterns they drive in reporter genes from *Dll_LE2* (A); *spi_LE2* (B); *noc_LE2* (C); *Antp_LE1* (D); *Antp_LE2* (E); *Ote/fj_LE* (F).

## References

1. Levine M. Transcriptional enhancers in animal development and evolution. Curr Biol. 2010;20(17):R754–63. doi: 10.1016/j.cub.2010.06.070. PubMed PMID: 20833320; PubMed Central PMCID: PMCPMC4280268.

2. Estella C, Voutev R, Mann RS. A dynamic network of morphogens and transcription factors patterns the fly leg. Curr Top Dev Biol. 2012;98:173–98. doi: 10.1016/B978-0-12-386499-4.00007-0. PubMed PMID: 22305163; PubMed Central PMCID: PMCPMC3918458.

3. Cohen SM. Specification of limb development in the Drosophila embryo by positional cues from segmentation genes. Nature. 1990;343(6254):173–7. Epub 1990/01/11. doi: 10.1038/343173a0. PubMed PMID: 2296309.

4. Estella C, Rieckhof G, Calleja M, Morata G. The role of buttonhead and Sp1 in the development of the ventral imaginal discs of Drosophila. Development. 2003;130(24):5929–41. Epub 2003/10/17. doi: 10.1242/dev.00832 dev.00832 [pii]. PubMed PMID: 14561634.

5. Estella C, Mann RS. Non-redundant selector and growth-promoting functions of two sister genes, buttonhead and Sp1, in Drosophila leg development. PLoS Genet. 2010;6(6):e1001001. Epub 2010/06/30. doi: 10.1371/journal.pgen.1001001. PubMed PMID: 20585625; PubMed Central PMCID: PMC2891808.

6. Campbell G, Weaver T, Tomlinson A. Axis specification in the developing Drosophila appendage: the role of wingless, decapentaplegic, and the homeobox gene aristaless. Cell. 1993;74(6):1113–23. PubMed PMID: 8104704.

7. Diaz-Benjumea FJ, Cohen B, Cohen SM. Cell interaction between compartments establishes the proximal-distal axis of Drosophila legs. Nature. 1994;372(6502):175–9. PubMed PMID: 7969450.

8. Lecuit T, Cohen SM. Proximal-distal axis formation in the Drosophila leg. Nature. 1997;388(6638):139–45. PubMed PMID: 9217152.

9. Estella C, McKay DJ, Mann RS. Molecular integration of wingless, decapentaplegic, and autoregulatory inputs into Distalless during Drosophila leg development. Dev Cell. 2008;14(1):86–96. PubMed PMID: 18194655.

10. Giorgianni MW, Mann RS. Establishment of medial fates along the proximodistal axis of the Drosophila leg through direct activation of dachshund by Distalless. Dev Cell. 2011;20(4):455–68. Epub 2011/04/19. doi: S1534-5807(11)00123-7 [pii] 10.1016/j.devcel.2011.03.017. PubMed PMID: 21497759; PubMed Central PMCID: PMC3087180.

11. Campbell G. Distalization of the Drosophila leg by graded EGF-receptor activity. Nature. 2002;418(6899):781–5. PubMed PMID: 12181568.

12. Galindo MI, Bishop SA, Greig S, Couso JP. Leg patterning driven by proximal-distal interactions and EGFR signaling. Science. 2002;297(5579):256–9. PubMed PMID: 12114628.

13. Shilo BZ. Signaling by the Drosophila epidermal growth factor receptor pathway during development. Exp Cell Res. 2003;284(1):140–9. PubMed PMID: 12648473.

14. Galindo MI, Bishop SA, Couso JP. Dynamic EGFR-Ras signalling in Drosophila leg development. Dev Dyn. 2005;233(4):1496–508. PubMed PMID: 15965980.

15. Campbell G. Regulation of gene expression in the distal region of the Drosophila leg by the Hox11 homolog, C15. Dev Biol. 2005;278(2):607–18. doi: 10.1016/j.ydbio.2004.12.009. PubMed PMID: 15680373.

16. Wasserman JD, Urban S, Freeman M. A family of rhomboid-like genes: Drosophila rhomboid-1 and roughoid/rhomboid-3 cooperate to activate EGF receptor signaling. Genes Dev. 2000;14(13):1651–63. PubMed PMID: 10887159.

17. Gratz SJ, Cummings AM, Nguyen JN, Hamm DC, Donohue LK, Harrison MM, et al. Genome engineering of Drosophila with the CRISPR RNA-guided Cas9 nuclease. Genetics. 2013;194(4):1029–35. doi: 10.1534/genetics.113.152710. PubMed PMID: 23709638; PubMed Central PMCID: PMCPMC3730909.

18. Port F, Chen HM, Lee T, Bullock SL. Optimized CRISPR/Cas tools for efficient germline and somatic genome engineering in Drosophila. Proc Natl Acad Sci U S A. 2014;111(29):E2967–76. Epub 2014/07/09. doi: 1405500111 [pii] 10.1073/pnas.1405500111. PubMed PMID: 25002478; PubMed Central PMCID: PMC4115528.

19. Kondo S, Ueda R. Highly improved gene targeting by germline-specific Cas9 expression in Drosophila. Genetics. 2013;195(3):715–21. Epub 2013/09/05. doi: genetics.113.156737 [pii] 10.1534/genetics.113.156737. PubMed PMID: 24002648; PubMed Central PMCID: PMC3813859.

20. Kojima T, Sato M, Saigo K. Formation and specification of distal leg segments in Drosophila by dual Bar homeobox genes, BarH1 and BarH2. Development. 2000;127(4):769–78. Epub 2000/01/29. PubMed PMID: 10648235.

21. Powell J. Progress and Prospects in Evolutionary Biology: The Drosophila Model: Oxford University Press; 1997 Sep 4, 1997.

22. McDonald JA, Pinheiro EM, Kadlec L, Schupbach T, Montell DJ. Multiple EGFR ligands participate in guiding migrating border cells. Dev Biol. 2006;296(1):94–103. doi: 10.1016/j.ydbio.2006.04.438. PubMed PMID: 16712835.

23. Lan L, Lin S, Zhang S, Cohen RS. Evidence for a transport-trap mode of Drosophila melanogaster gurken mRNA localization. PLoS One. 2010;5(11):e15448. doi: 10.1371/journal.pone.0015448. PubMed PMID: 21103393; PubMed Central PMCID: PMCPMC2980492.

24. Simcox AA, Grumbling G, Schnepp B, Bennington-Mathias C, Hersperger E, Shearn A. Molecular, phenotypic, and expression analysis of vein, a gene required for growth of the Drosophila wing disc. Dev Biol. 1996;177(2):475–89. doi: 10.1006/dbio.1996.0179. PubMed PMID: 8806825.

25. Donaldson T, Wang SH, Jacobsen TL, Schnepp B, Price J, Simcox A. Regulation of the Drosophila epidermal growth factor-ligand vein is mediated by multiple domains. Genetics. 2004;167(2):687–98. doi: 10.1534/genetics.103.019588. PubMed PMID: 15238521; PubMed Central PMCID: PMCPMC1470887.

26. Yu L, Lee T, Lin N, Wolf MJ. Affecting Rhomboid-3 function causes a dilated heart in adult Drosophila. PLoS Genet. 2010;6(5):e1000969. doi: 10.1371/journal.pgen.1000969. PubMed PMID: 20523889; PubMed Central PMCID: PMCPMC2877733.

27. Tchankouo-Nguetcheu S, Udinotti M, Durand M, Meng TC, Taouis M, Rabinow L. Negative regulation of MAP kinase signaling in Drosophila by Ptp61F/PTP1B. Mol Genet Genomics. 2014;289(5):795–806. doi: 10.1007/s00438-014-0852-2. PubMed PMID: 24752400.

28. Gallio M, Englund C, Kylsten P, Samakovlis C. Rhomboid 3 orchestrates Slit-independent repulsion of tracheal branches at the CNS midline. Development. 2004;131(15):3605–14. doi: 10.1242/dev.01242. PubMed PMID: 15229181.

29. Cohen SM, Bronner G, Kuttner F, Jurgens G, Jackle H. Distal-less encodes a homoeodomain protein required for limb development in Drosophila. Nature. 1989;338(6214):432–4. PubMed PMID: 2564639.

30. Scholz H, Deatrick J, Klaes A, Klambt C. Genetic dissection of pointed, a Drosophila gene encoding two ETS-related proteins. Genetics. 1993;135(2):455–68. PubMed PMID: 8244007.

31. Kumar JP, Tio M, Hsiung F, Akopyan S, Gabay L, Seger R, et al. Dissecting the roles of the Drosophila EGF receptor in eye development and MAP kinase activation. Development. 1998;125(19):3875–85. PubMed PMID: 9729495.

32. Roch F, Jimenez G, Casanova J. EGFR signalling inhibits Capicua-dependent repression during specification of Drosophila wing veins. Development. 2002;129(4):993–1002. PubMed PMID: 11861482.

33. Lee T, Luo L. Mosaic analysis with a repressible cell marker (MARCM) for Drosophila neural development. Trends Neurosci. 2001;24(5):251–4. PubMed PMID: 11311363.

34. Struhl G, Basler K. Organizing activity of wingless protein in Drosophila. Cell. 1993;72(4):527–40. Epub 1993/02/26. doi: 0092-8674(93)90072-X [pii]. PubMed PMID: 8440019.

35. Zhu LJ, Christensen RG, Kazemian M, Hull CJ, Enuameh MS, Basciotta MD, et al. FlyFactorSurvey: a database of Drosophila transcription factor binding specificities determined using the bacterial one-hybrid system. Nucleic Acids Res. 2011;39(Database issue):D111–7. doi: 10.1093/nar/gkq858. PubMed PMID: 21097781; PubMed Central PMCID: PMCPMC3013762.

36. Sosinsky A, Bonin CP, Mann RS, Honig B. Target Explorer: An automated tool for the identification of new target genes for a specified set of transcription factors. Nucleic Acids Res. 2003;31(13):3589–92. PubMed PMID: 12824372.

37. McKay DJ, Lieb JD. A common set of DNA regulatory elements shapes Drosophila appendages. Dev Cell. 2013;27(3):306–18. doi: 10.1016/j.devcel.2013.10.009. PubMed PMID: 24229644; PubMed Central PMCID: PMCPMC3866527.

38. McKay DJ, Estella C, Mann RS. The origins of the Drosophila leg revealed by the cis-regulatory architecture of the Distalless gene. Development. 2009;136(1):61–71. PubMed PMID: 19036798.

39. Galindo MI, Fernandez-Garza D, Phillips R, Couso JP. Control of Distal-less expression in the Drosophila appendages by functional 3’ enhancers. Dev Biol. 2011;353(2):396–410. Epub 2011/02/16. doi: S0012-1606(11)00092-3 [pii] 10.1016/j.ydbio.2011.02.005. PubMed PMID: 21320482.

40. Jory A, Estella C, Giorgianni MW, Slattery M, Laverty TR, Rubin GM, et al. A survey of 6,300 genomic fragments for cis-regulatory activity in the imaginal discs of Drosophila melanogaster. Cell Rep. 2012;2(4):1014–24. doi: 10.1016/j.celrep.2012.09.010. PubMed PMID: 23063361; PubMed Central PMCID: PMCPMC3483442.

41. Wee P, Wang Z. Epidermal Growth Factor Receptor Cell Proliferation Signaling Pathways. Cancers (Basel). 2017;9(5). doi: 10.3390/cancers9050052. PubMed PMID: 28513565; PubMed Central PMCID: PMCPMC5447962.

42. Curtiss J, Halder G, Mlodzik M. Selector and signalling molecules cooperate in organ patterning. Nat Cell Biol. 2002;4(3):E48–51. doi: 10.1038/ncb0302-e48. PubMed PMID: 11875444.

43. Roberts PJ, Der CJ. Targeting the Raf-MEK-ERK mitogen-activated protein kinase cascade for the treatment of cancer. Oncogene. 2007;26(22):3291–310. PubMed PMID: 17496923.

44. Chaleshi V, Haghighi MM, Savabkar S, Zali N, Vahedi M, Khanyaghma M, et al. Correlation between the EGF gene intronic polymorphism, rs2298979, and colorectal cancer. Oncol Lett. 2013;6(4):1079–83. doi: 10.3892/ol.2013.1481. PubMed PMID: 24137467; PubMed Central PMCID: PMCPMC3796412.

45. Chen X, Yang G, Zhang D, Zhang W, Zou H, Zhao H, et al. Association between the epidermal growth factor +61 G/A polymorphism and glioma risk: a meta-analysis. PLoS One. 2014;9(4):e95139. doi: 10.1371/journal.pone.0095139. PubMed PMID: 24740103; PubMed Central PMCID: PMCPMC3989292.

46. Kristensen VN, Edvardsen H, Tsalenko A, Nordgard SH, Sorlie T, Sharan R, et al. Genetic variation in putative regulatory loci controlling gene expression in breast cancer. Proc Natl Acad Sci U S A. 2006;103(20):7735–40. doi: 10.1073/pnas.0601893103. PubMed PMID: 16684880; PubMed Central PMCID: PMCPMC1458617.

47. Cordoba S, Requena D, Jory A, Saiz A, Estella C. The evolutionarily conserved transcription factor Sp1 controls appendage growth through Notch signaling. Development. 2016;143(19):3623–31. doi: 10.1242/dev.138735. PubMed PMID: 27578786.

48. Austin CL, Manivannan SN, Simcox A. TGF-alpha ligands can substitute for the neuregulin Vein in Drosophila development. Development. 2014;141(21):4110–4. doi: 10.1242/dev.110171. PubMed PMID: 25336739.

49. Slattery M, Voutev R, Ma L, Negre N, White KP, Mann RS. Divergent transcriptional regulatory logic at the intersection of tissue growth and developmental patterning. PLoS Genet. 2013;9(9):e1003753. doi: 10.1371/journal.pgen.1003753. PubMed PMID: 24039600; PubMed Central PMCID: PMCPMC3764184.

50. Bischof J, Maeda RK, Hediger M, Karch F, Basler K. An optimized transgenesis system for Drosophila using germ-line-specific phiC31 integrases. Proc Natl Acad Sci U S A. 2007;104(9):3312–7. doi: 10.1073/pnas.0611511104. PubMed PMID: 17360644; PubMed Central PMCID: PMCPMC1805588.

51. Groth AC, Fish M, Nusse R, Calos MP. Construction of transgenic Drosophila by using the site-specific integrase from phage phiC31. Genetics. 2004;166(4):1775–82. PubMed PMID: 15126397.

52. Voutev R, Mann RS. Robust PhiC31-Mediated Genome Engineering in Drosophila melanogaster Using Minimal attP/attB Phage Sites. G3 (Bethesda). 2018. doi: 10.1534/g3.118.200051. PubMed PMID: 29523637.

53. Gebelein B, Culi J, Ryoo HD, Zhang W, Mann RS. Specificity of Distalless repression and limb primordia development by abdominal Hox proteins. Dev Cell. 2002;3(4):487–98. Epub 2002/11/01. doi: S1534580702002575 [pii]. PubMed PMID: 12408801.

54. Noro B, Culi J, McKay DJ, Zhang W, Mann RS. Distinct functions of homeodomain-containing and homeodomain-less isoforms encoded by homothorax. Genes Dev. 2006;20(12):1636–50. doi: 10.1101/gad.1412606. PubMed PMID: 16778079; PubMed Central PMCID: PMCPMC1482483.

55. Charlton-Perkins M, Whitaker SL, Fei Y, Xie B, Li-Kroeger D, Gebelein B, et al. Prospero and Pax2 combinatorially control neural cell fate decisions by modulating Ras- and Notch-dependent signaling. Neural Dev. 2011;6:20. doi: 10.1186/1749-8104-6-20. PubMed PMID: 21539742; PubMed Central PMCID: PMCPMC3123624.

56. Higashijima S, Michiue T, Emori Y, Saigo K. Subtype determination of Drosophila embryonic external sensory organs by redundant homeo box genes BarH1 and BarH2. Genes Dev. 1992;6(6):1005–18. PubMed PMID: 1350558.

57. Mardon G, Solomon NM, Rubin GM. dachshund encodes a nuclear protein required for normal eye and leg development in Drosophila. Development. 1994;120(12):3473–86. PubMed PMID: 7821215.

58. Morris CA, Benson E, White-Cooper H. Determination of gene expression patterns using in situ hybridization to Drosophila testes. Nat Protoc. 2009;4(12):1807–19. doi: 10.1038/nprot.2009.192. PubMed PMID: 20010932.

59. Slattery M, Ma L, Negre N, White KP, Mann RS. Genome-wide tissue-specific occupancy of the Hox protein Ultrabithorax and Hox cofactor Homothorax in Drosophila. PLoS One. 2011;6(4):e14686. doi: 10.1371/journal.pone.0014686. PubMed PMID: 21483663; PubMed Central PMCID: PMCPMC3071676.

60. Li H, Durbin R. Fast and accurate short read alignment with Burrows-Wheeler transform. Bioinformatics. 2009;25(14):1754–60. doi: 10.1093/bioinformatics/btp324. PubMed PMID: 19451168; PubMed Central PMCID: PMCPMC2705234.

61. Zhang Y, Liu T, Meyer CA, Eeckhoute J, Johnson DS, Bernstein BE, et al. Model-based analysis of ChIP-Seq (MACS). Genome Biol. 2008;9(9):R137. doi: 10.1186/gb-2008-9-9-r137. PubMed PMID: 18798982; PubMed Central PMCID: PMCPMC2592715.

